# Giardia Antagonizes Beneficial Functions of Indigenous and Therapeutic Intestinal Bacteria during Malnutrition

**DOI:** 10.1101/2024.01.22.575921

**Authors:** Aadra P. Bhatt, Jason W. Arnold, Muyiwa Awoniyi, Shan Sun, Verônica Feijoli Santiago, Pedro Henrique Quintela, Kenneth Walsh, Renay Ngobeni, Brenna Hansen, Ajay Gulati, Ian M. Carroll, M. Andrea Azcarate-Peril, Anthony A. Fodor, Jonathan Swann, Luther A. Bartelt

## Abstract

Undernutrition in children commonly disrupts the structure and function of the small intestinal microbial community, leading to enteropathies, compromised metabolic health, and impaired growth and development. The mechanisms by which diet and microbes mediate the balance between commensal and pathogenic intestinal flora remain elusive. In a murine model of undernutrition, we investigated the direct interactions *Giardia lamblia,* a prevalent small intestinal pathogen, on indigenous microbiota and specifically on Lactobacillus strains known for their mucosal and growth homeostatic properties. Our research reveals that *Giardia* colonization shifts the balance of lactic acid bacteria, causing a relative decrease in *Lactobacillus spp*. and an increase in *Bifidobacterium spp*. This alteration corresponds with a decrease in multiple indicators of mucosal and nutritional homeostasis. Additionally, protein-deficient conditions coupled with *Giardia* infection exacerbate the rise of primary bile acids and susceptibility to bile acid-induced intestinal barrier damage. In epithelial cell monolayers, *Lactobacillus spp*. mitigated bile acid-induced permeability, showing strain-dependent protective effects. *In vivo, L. plantarum,* either alone or within a *Lactobacillus* spp consortium, facilitated growth in protein-deficient mice, an effect attenuated by *Giardia*, despite not inhibiting Lactobacillus colonization. These results highlight Giardia’s potential role as a disruptor of probiotic functional activity, underscoring the imperative for further research into the complex interactions between parasites and bacteria under conditions of nutritional deficiency.

## INTRODUCTION

Childhood undernutrition and linear growth impairment are a widespread and complex global health problem^1^ ^2^. Beyond a state of nutritional deficiency, our current understanding suggests that childhood undernutrition also includes nutrient-dependent disruption in the absorptive and metabolic functions of the developing small intestine (SI) ^3^. These functions are known to be influenced by the density, composition, and function of resident intestinal microbes. Although nutrient absorption predominantly occurs in the SI, most studies examining the influences of intestinal microbiota on host nutritional and metabolic homeostasis utilize platform technologies that primarily profile colonic microbial communities. Recent microbial community profiling of duodenal aspirates and biopsies collected from undernourished children unresponsive to nutritional supplementation revealed a shift towards predominantly oral-mucosal bacteria ^4, 5^ and notably a reduction in prototypic SI commensals like *Lactobacillus spp*.^6^. Additionally, conventional intestinal pathogens, such as diarrheagenic *Escherichia coli* types, *Campylobacter spp.* and *Giardia lamblia* that independently associate with impaired childhood growth ^7^ have also been detected in SI communities from treatment-refractory undernourished children ^5, 8^. The potential consequences of interactions between the co-occurrence of these conventional gut pathogens and altered SI microbial communities in children with undernutrition is poorly understood.

*G. lamblia* (*Giardia*) is implicated in several studies examining the consequences of disrupted SI microbial communities on childhood growth and intestinal function. We and others have shown that *Giardia* may restrict child linear growth through dose-dependent SI epithelial cell permeability dysfunction and disruptions in microbial-host nutrient homeostasis ^9^ ^10^. *Giardia* also associates with markers of SI bacterial overgrowth ^11^ and diminished markers of lymphocyte activation ^8^. Using gnotobiotic wild-type mice, we have also shown that *Giardia*-mediated growth impairment results from the convergence of two independent factors: inadequate protein intake, and perturbed intestinal microbiota ^10^ ^9^. These data link microbiota-host homeostasis with the presence of *Giardia.* However, the directionality of this interaction, which specific bacteria are most relevant, and the consequences of these disruptions on intestinal development and overall growth trajectories are not well understood.

Trials to remediate disrupted intestinal microbiota and restore healthy child growth are underway (e.g. NCT05570045, NCT00118872). Among the different strategies, probiotic interventions with defined bacteria, like certain *Lactobacillus spp.* strains have been appealing. In animal models, specific *Lactobacillus spp.* probiotics have been shown to support linear growth in mono-associated undernourished mice ^12^, protect intestinal epithelial cell integrity and support epithelial repair ^13, 14^, and regulate mucosal and immune responses to pathogens like *Giardia*^15–17^. However, undernutrition can lead to rapid metabolic adaptations in *Lactobacillus spp.* and potential diminished mucosal-associated compartmentalization ^18^ that could limit beneficial host interactions from commensal and/or probiotic strains.

Here we investigated interactions between *Giardia* and specific *Lactobacillus spp.* in the nourished and undernourished murine host. We find that during protein deprivation, *Giardia* contributes to specific reductions in specific *Lactobacillus* species abundance, relative to other commensal bacteria. These changes are coupled with decreases in indicators of disrupted intestinal mucosal and nutrient homeostasis. We find that protein undernutrition increases intestinal primary bile acids, and the additional presence of *Giardia* associates with indicators of increased bile acid-associated intestinal epithelial cell (IEC) injury. In an *in vitro* model of IEC monolayers, bile acids cause permeability defects at physiological concentrations, but a bile salt hydrolase-expressing *L. plantarum* WCSF1 protects against this injury. While a consortium of commensal *Lactobacillus spp.* or mono-association with *L. plantarum* WCSF1 supports weight gain in gnotobiotic protein deprived mice, co-colonization with *Giardia* inhibits this effect. These findings indicate that *Giardia* colonization may contribute to loss of microbiota-host homeostasis through loss of commensal *Lactobacillus* functions. These findings implicate an important role for *Giardia* modulation of small intestinal microbiota composition, particularly in the context of engraftment and/or efficacy of bacterial-based therapies and live biotherapeutic products (LBPs) used to correct malnutrition.

## RESULTS

### *Giardia* diminishes *Lactobacillus* and increases *Bifidobacterium* relative abundances in protein-deprived mice

We previously reported a specific pathogen free (SPF) model of juvenile protein-deprived mice that develop *Giardia*-enteropathy^19^. In the present study, we profiled intestinal bacterial communities using fecal 16S rRNA amplicon sequencing between 9-11 days following cyst challenge, a timepoint at which *Giardia*-colonized **p**rotein-deficient **d**iet (**PD**)-fed mice have reproducibly established growth restriction and increased intestinal permeability ^9, 10^. While protein deficiency drove fecal microbiota community changes, we found that PD-fed mice challenged with *Giardia* had a further alteration in community profiles, compared to those challenged with control (PBS) (**Fig. 1A**). In PD-fed mice, the presence of Giardia resulted in genus-level alterations in microbiota composition, characterized by a reduction in *Lactobacillus,* and a reciprocal increase in *Bifidobacterium* (**Fig. 1B**). Targeted qPCR using feces from PD-fed *Giardia*-challenged mice confirmed a >5-fold reduction in *Lactobacillus* (P<0.01) and a >2-fold increase in *Bifidobacterium* (P<0.05) relative to total bacteria, as estimated by universal 16S qPCR (**Fig. 1C**). Within the *Giardia* trophozoite-rich duodenum of PD-fed mice, there was a ∼4-fold decrease in *Lactobacillus* compared with diet matched controls (P<0.01). However, *Bifidobacterium* in the duodenum did not appreciably (P=0.132) differ in the presence or absence of *Giardia* (**Fig. 1D**), indicating that protein deficiency may be a primary driver of *Bifidobacterium* abundance.

**Figure 1.**
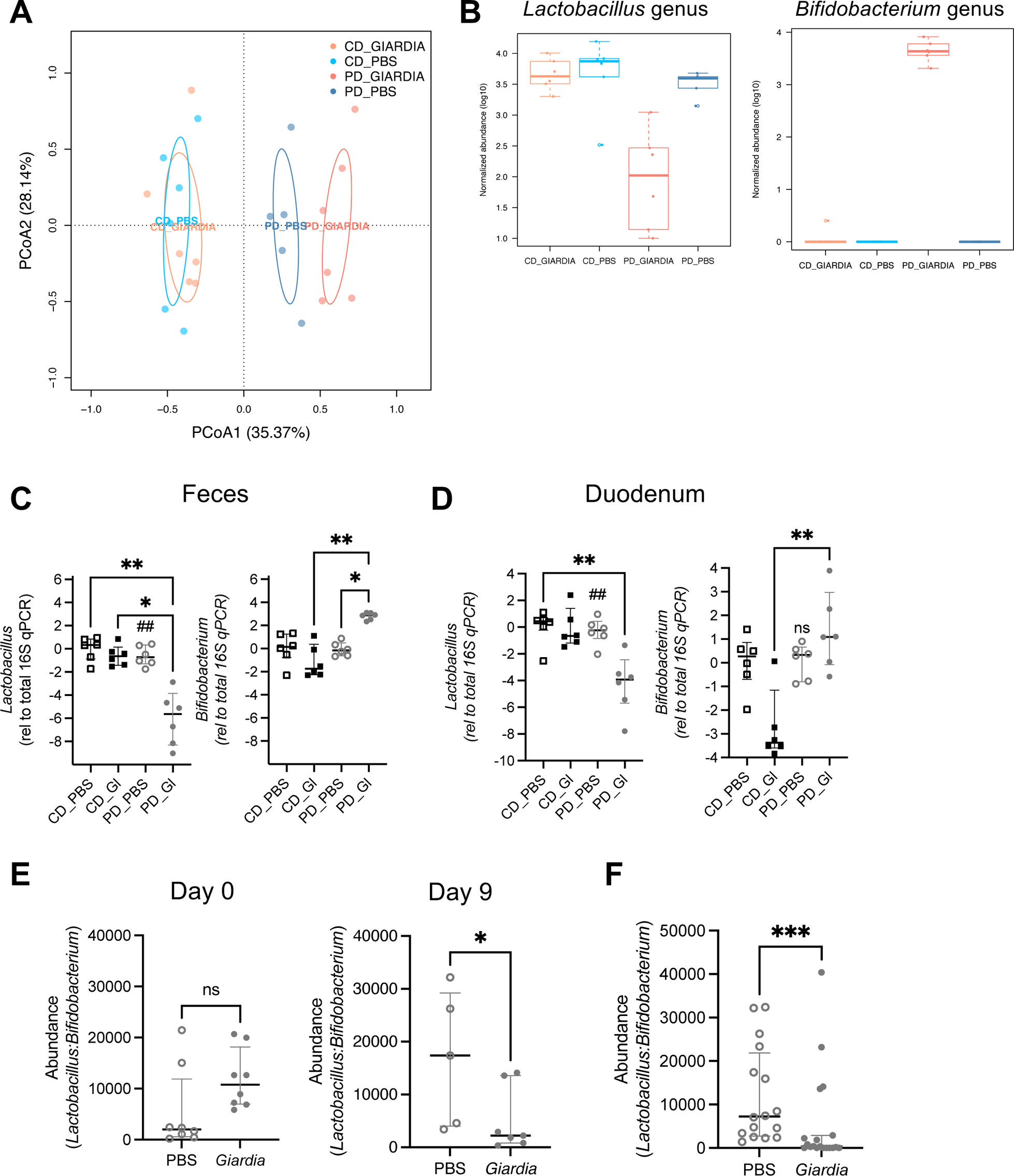
*Giardia* differentially alters *Lactobacillus* and *Bifidobacterium* abundances in protein-deprived mice. **(A)** PCoA plots of fecal taxonomic profiles at genus level from mice fed either a control diet (CD) or an isocaloric protein deficient diet (PD). Samples were collected 10 days after challenge with 10^5^ *G. lamblia* cysts or PBS as indicated (N=6 mice per group, N=2-3 cages per group). PD_PBS vs PD_Giardia: **R2 = 0.23621 P= 0.037;** CD_PBS vs CD_Giardia: **R2 = 0.02552 P= 0.907;** CD_PBS vs PD_PBS: **R2 = 0.22498 P= 0.05;** CD_Giardia vs PD_Giardia: **R2 = 0.35802 P= 0.002** (PERMANOVA test with 999 permutations). **(B)** Normalized abundances (log10, median ± IQR) of genus-level assignments for *Lactobacillus* (left) or *Bifidobacterium* (right). *P<0.05, Mann-Whitney U-test. **(C, D)** Abundances of *Lactobacillus* and *Bifidobacterium* by qPCR from paired fecal **(C)** and **(D)** duodenum samples as labeled. Data are represented as relative to total 16S qPCR. *P<0.05, **P<0.01, Kruskal-Wallis with Dunn’s test for multiple comparisons as indicated, N=6 per group. **(E)** Ratios of relative abundances of *Lactobacillus:Bifidobacterium* in fecal samples from PD-diet fed *Giardia* challenged or PBS control mice aggregated day 9-11 after *Giardia* challenge from three independent experiments, N=16-19 per group. ***P<0.001 (Mann-Whitney U-test, median ± IQR, N=6 per group). **(F)** Ratios of abundances of *Lactobacillus:Bifidobacterium* in fecal samples from PD-fed, PBS- or *Giardia*-challenged mice from individual representative experiment: Day of *Giardia* challenge (day 0, **Left**) and Day 9 after *Giardia* challenge (**Right)**,. *P<0.5 (Mann-Whitney U-test, median ± IQR, N=5-8 per group.

We individually housed mice and used genus-specific and universal 16S qPCR to quantify baseline *Lactobacillus* and *Bifidobacterium* to account for potential confounders arising from their differences prior to *Giardia* exposure. In this experiment, baseline *Lactobacillus:Bifidobacterium* relative abundances were unequally distributed with slightly higher ratios in the *Giardia*-challenged mice. However, by 9 days after *Giardia* challenge, *Lactobacillus:Bifidobacterium* relative abundances had significantly decreased compared with PBS-controls (**Fig. 1E**). This decrease was observed in additional age- and diet-matched independent repeats of this experiment, albeit with varying ratios, likely resulting from cohousing (**Supplemental Fig. 1A, 1B**). Aggregating all age- and diet-matched experiments, we found that the presence of *Giardia* reproducibly and significantly diminished *Lactobacillus:Bifidobacterium* ratios within 9-11 days post colonization (**Fig. 1F**).

### *Giardia* infection diminishes distinct markers of epithelial cell homeostasis

Given the *Giardia*-mediated reduction in *Lactobacillus* abundance in the upper SI (**Fig. 1B-F**), we next tested whether epithelial homeostasis markers associated with commensal *Lactobacillus* functions were also diminished in *Giardia*-challenged PD-diet fed mice. These include markers of intestinal mucosal defense responses ^20^, regeneration and repair ^14^, and regulation of transcellular protein transport ^21^, all reportedly modulated by *Lactobacillus*. First, compared with PBS-challenged PD-fed controls, *Giardia*-challenged animals had reduced SI gene expression of innate responses regulated by RegIIIγ and IL22 in response to Gram-positive bacteria (**Fig. 2A**). Interestingly, we did not observe upregulation in matrix metalloprotease 7 (MMP7) following *Giardia* challenge, which has previously been reported to occur following *Giardia lamblia* infection during adequate nourishment, and important for controlling parasite numbers ^22^. Compensatory crypt expansion seen in nourished *Giardia*-challenged mice (**Fig. 2C**) was not observed in PD-fed *Giardia*-challenged mice (**Fig. 2B**). Reflecting previous findings in malnourished rats ^23^, we found that protein deprivation led to compensatory upregulation of the oligopeptide transporter *PepT1,* which was partially diminished by the presence of *Giardia* (**Fig. 2D**).

**Figure 2.**
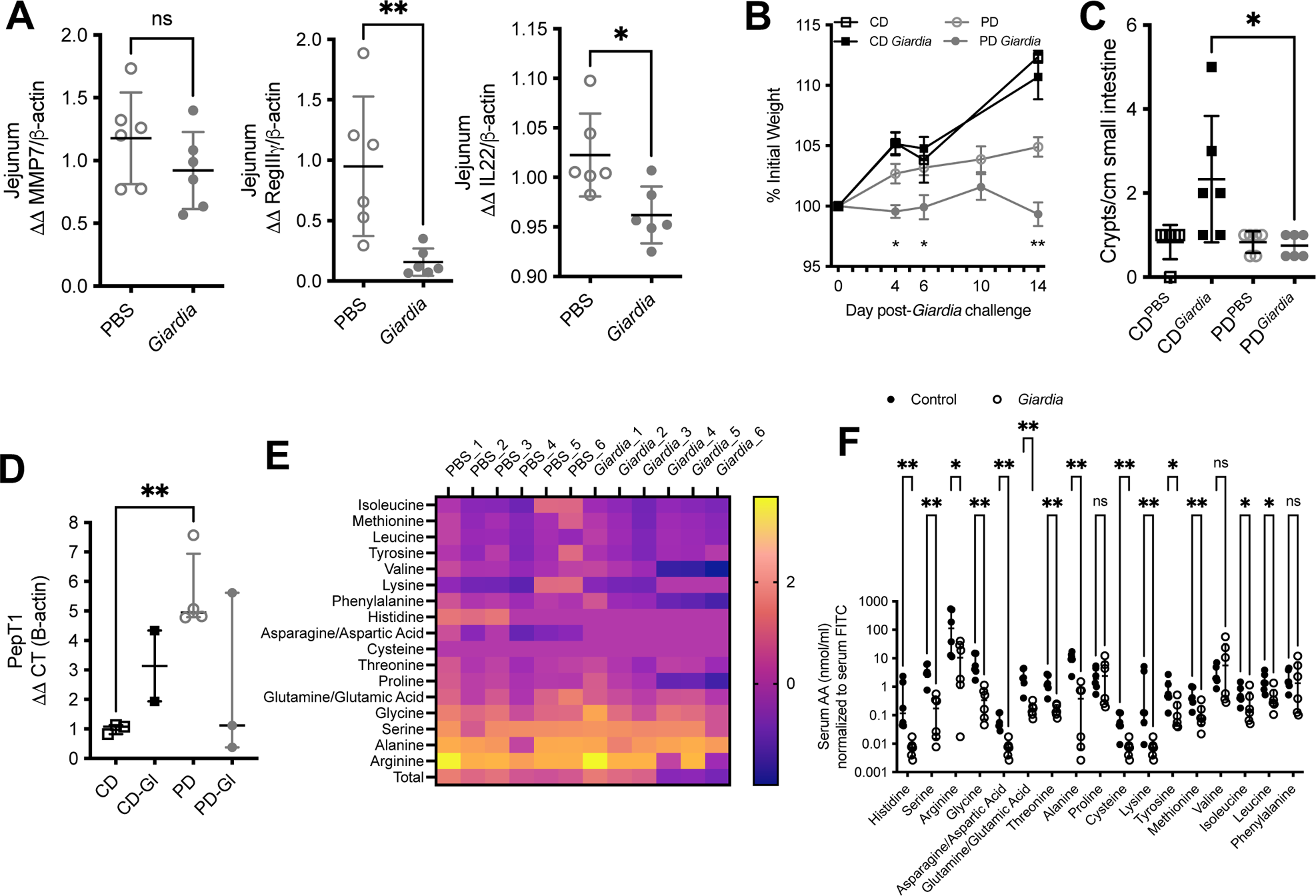
*Giardia* alters markers of intestinal epithelial cell homeostasis that are associated with commensal bacteria functions. **A**-**C)** Relative gene expression of innate mucosal responses in jejunum of PD-fed PBS controls and 10 days after *Giardia* challenge as indicated: **(A)** *RegIII*γ*,* **(B)** *MMP7,* **(C)** *IL22*. *P<0.05 and **P<0.01, Mann-Whitney test, N=6 per group. **(D)** Relative expression of oligopeptide transporter *PepT1* in duodenum of CD or PD-diet fed PBS controls and 10 days after *Giardia* challenge. *P<0.05, Mann-Whitney, N=3 per group. For A-D, data are shown as ΔΔ to β-actin housekeeping gene and normalized to PBS controls (median ± IQR). **(B)** Growth as % initial weight of CD or PD-fed PBS controls or *Giardia*-challenged mice. Three-week-old specific pathogen free (SPF) mice were acclimated to respective diets for 10 days prior to challenge with 10^5^ *G. lamblia* cysts or PBS. *P<0.05, ****P<0.0001 Two-way ANOVA with Tukey’s post hoc analysis for PD-*Giardia* vs. PD (mean ± SEM, N=6 per group). **(C)** Enumeration of crypts isolated from the small intestine of mice in Figure E. *P<0.05, Kruskal-Wallis with Dunn’s test for multiple comparisons as indicated, N=6 per group (median ± IQR). **(E)** Heatmap representation of ratio of free amino acids measured in serum to fecal compartments [log transformed] in PD-diet fed PBS controls and 10 days after *Giardia* challenge. **(F)** free amino acids in serum normalized to concomitant measurement of serum FITC as a measure of intestinal permeability. *P<0.05, **P<0.01, Multiple t-tests, Benjamini, Krieger, and Yekutielie 2-step method with FDR 5% (median ± IQR, N=6 per group).

In accordance with the published literature, we found that protein deficiency significantly altered serum (**Supplemental Fig. 2A-C**) and fecal (**Supplemental Fig. 2D-E**) pools of free amino acids, regardless of *Giardia* infection status. Serum: fecal ratios of free amino acids in *Giardia*-challenged PD-diet fed mice tended to be lower than PBS controls (**Fig. 2E, Supp. Fig. 2G**). We accounted for unregulated paracellular amino acid flux reported to occur with increased gut permeability^9^ by measuring serum FITC-dextran. Adjusting for permeability, *Giardia* challenge significantly reduced serum histidine, serine, arginine, glycine, asparagine/aspartic acid, glutamine/glutamic acid, threonine, alanine, cysteine, lysine, tyrosine, methionine, isoleucine, and leucine (**Fig. 2F**). Levels of proline, valine, and phenylalanine were similar in both groups. These data demonstrate that *Giardia* infection exacerbates the already disrupted protein absorption occurring in protein malnourishment thus further reducing circulating amino acids.

Together these findings indicate that *Giardia* colonization in protein undernutrition impedes compensatory mucosal and nutrient responses to restore homeostasis, which rely upon function of critical commensal bacteria which are lost due to parasite infection.

### Protein deficiency increases primary bile acids, which positively correlates with *Giardia*-mediated barrier defects

A primary metabolic function of several commensal *Lactobacillus* species is regulation of bile acid homeostasis through the expression of bile salt hydrolases (*bsh*) that deconjugate glyco- or tauro-conjugated primary bile acids ^24^. Allochthonous *Lactobacillus* strains diminish transient *Giardia* infection in murine models through this deconjugation activity, increasing the pool of deconjugated bile salts ^25–27^. In a similar vein, adult mice persistently infected with *G. lamblia* had decreased amounts of taurine-conjugated bile acids, and increased levels of the corresponding unconjugated bile acids ^28^. To determine if altered *Lactobacillus:Bifidobacterium* ratios contributed to disrupted bile acid homeostasis during protein deprivation, we profiled fecal and serum bile acids at 10/11 days post-*Giardia* challenge in PD-fed and adequately nourished mice (20 days total on either PD or regular chow). Fecal bile acid profiles were largely diet-dependent (**Fig. 3A, C**), without a remarkable influence of *Giardia* challenge (**Supplemental Fig. 3A, B**). Regardless of *Giardia* status, the PD-diet led to an increase of primary bile acids relative to secondary bile acids (**Fig. 3D-F, Supplemental Fig. 3B**). Comparisons between conjugated and unconjugated bile acids showed more variability between PD-diet fed mice (**Fig. 3C, 3D),** but overall diminished amounts of both glyco- and tauro-conjugated bile acids in PD-fed mice indicated preserved bacterial *bsh* activity (**Fig. 3C, D**).

**Figure 3.**
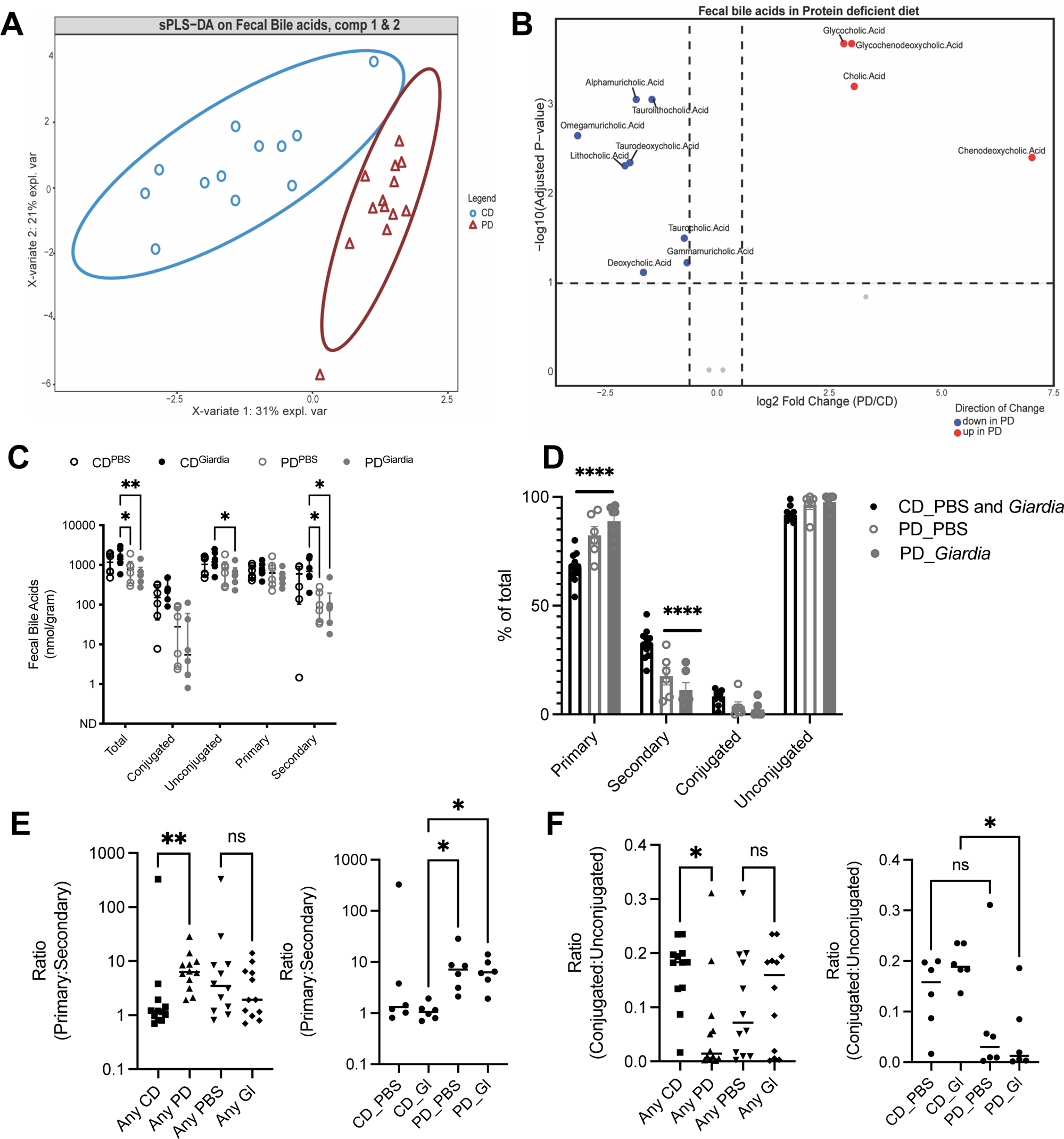
Protein deficiency restricts conversion of intestinal primary to secondary bile acids regardless of *Giardia* challenge. **(A)** sPLS-DA of faecal bile acids from mice fed a protein-deficient diet compared with the control diet (CD). **(B)** Volcano plot displays the significant bile acids in fecal samples with PD diet compared to control diet (CD). Taurine-conjugated (Taurolithocholic acid, Taurocholic acid, and Taurodeoxycholic acid) and secondary bile acids (Alphamuricholic acid, Omegamuricholic acid, Gammamuricholic acd, Lithocholic acid, and Deoxycholic acid) were identified as underrepresented in protein deficient diet (blue dots). In contrast, glycine-conjugated (Glycocholic acid, Glycochenodeoxycholic acid) and primary bile acids (Cholic acid and Chenodeoxycholic acid) were overrepresented in protein deficient diet (red dots) ((Wilcoxon rank-sum Test, 10% FDR). **(C)** Comparisons between total, conjugated, unconjugated, primary and secondary bile acids. *P<0.05, **P<0.01, Two-Way ANOVA with Tukey’s post-test analysis for multiple comparisons (median ± IQR, N=6 per group). **(D**) Percentage of total bile acids represented by primary, secondary, conjugated and unconjugated types. ****P<0.001 for PD_PBS or PD_*Giardia* vs. CD_PBS and *Giardia* groups combined. Two-Way ANOVA with Tukey’s post-test analysis for multiple comparisons (median ± IQR, N=6-12 per group). **(E**) Ratios of primary:secondary bile acids in aggregate groups by dietary or *Giardia* exposure (left) and individual groups (right). *P<0.05, **P<0.01 for indicated groups. Kruskal-Wallis with Dunn’s test for multiple comparisons as indicated, (median ± IQR, N=6 per group). **(F)** Ratios of conjugated:unconjugated bile acids in aggregate groups by dietary or *Giardia* exposure (left) and individual groups (right). *P<0.05 for indicated groups. Kruskal-Wallis with Dunn’s test for multiple comparisons as indicated, (median ± IQR, N=6 per group).

In our model, differences in serum bile acid signatures between groups were also primarily driven by diet (**Fig. 4A, B**). PD-diet increased total bile acids, with minimal impact of *Giardia* even in PD-fed mice (**Supplemental Fig. 4A, B**). Additionally, ileal expression of *Fxr* or *Tgr5* bile acid homeostasis regulators were comparable between groups (**Supplemental Figure 4C**). Although alterations in serum BA profiles are primarily driven by diet (**Fig. 4A, C),** serum chenodeoxycholic acid concentrations were significantly and positively correlated with FITC-dextran, a measure of intestinal permeability, in PD-fed *Giardia*-infected mice, indicating a positive association with *Giardia*-induced barrier defects (**Fig. 4D**). This relationship was absent in PD-fed PBS-challenged controls. These data suggest that although *Giardia* does not change global intestinal and serum bile acid profiles in our model, *Giardia* colonization may have altered host susceptibility to bile acid-induced epithelial barrier defects occurring during undernutrition ^29^ ^30^.

**Figure 4.**
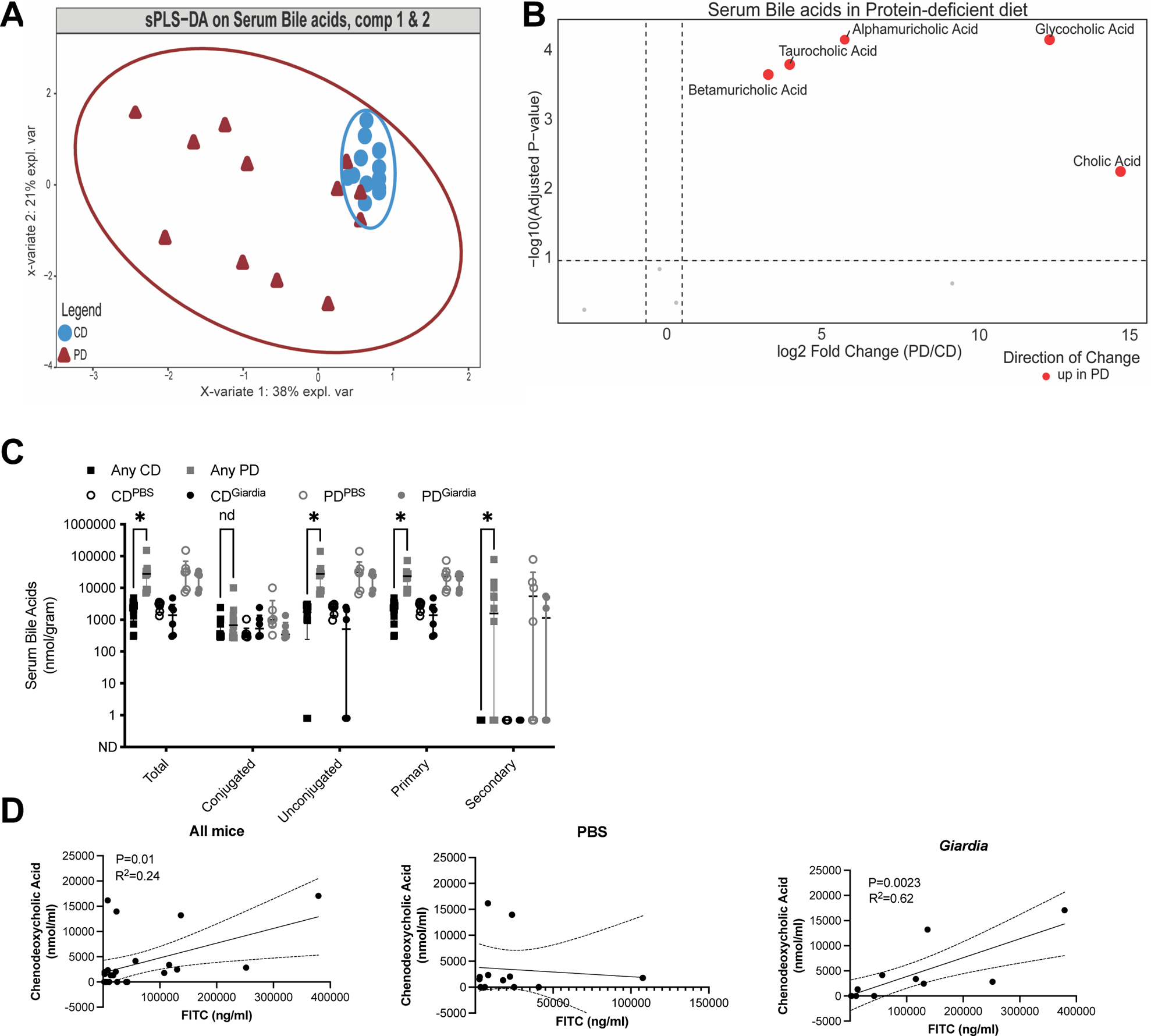
Protein deficiency increases circulating bile acids including chenodeoxycholic acid which correlates with severity of intestinal permeability in *Giardia* challenged mice. **(A**) sPLS-DA analysis of serum bile acids in mice fed a protein-deficient diet (PD) compared to mice fed a 20% protein diet (CD). **(B):** Volcano plot showing the significant abundance of serum bile acids in PD diet (Wilcoxon rank-sum Test, 10% FDR) with increased levels of glycine-conjugated bile acids (Glycocholic acid), Taurine-conjugated bile acids (Taurocholic acid), and secondary bile acids (Alphamuricholic acid and Betamuricholic acid), (red dots in the volcano plot). **(C)** Comparisons between total, conjugated, unconjugated, primary and secondary bile acids. *P<0.05, Multiple t-tests, Benjamini, Krieger, and Yekutielie 2-step method with FDR 5% (median ± IQR, N=6 per group). **(D**) Correlation between serum FITC (ng/mL) and chenodeoxycholic acid (nmol/mL) in PD-diet fed mice (left, all mice; middle just PBS controls; right just *Giardia* challenged mice). Simple linear regression (N=6 per group), R^2^=0.009238.

### Bile acid-induced barrier disruption is rescued by *bsh*-producing *L. plantarum*

Bile acids may have either homeostatic or toxic consequences on epithelial cells^31^ depending on the concentration and hydrophobicity of bile acid species that are present. We directly tested how a physiologically relevant mixture of bile acids perturbs intestinal epithelial cell permeability using the T84 cell line. Confluent T84 cell monolayers were grown until their transepithelial electrical resistance (TER) plateaued, indicative of a robust impermeant barrier. At this point, monolayers were exposed to a mixture of physiological bile acids dissolved in DMEM (termed pBA), or in *Giardia*-trophozoite supportive TYI-S-33 media; both contain 1%(w/v) bile acids in the proportions indicated in **Fig. 5A and Supplemental Fig. 5A**. Application of bile acids dissolved in either pBA or TYI-S-33 media resulted in immediate (within 2 hours) and sustained loss of TER indicative of a loss of barrier function (**Fig. 5B**). TER reduction occurred in a dose-dependent manner and had not recovered up to 12 hours following bile acid exposure.

**Figure 5.**
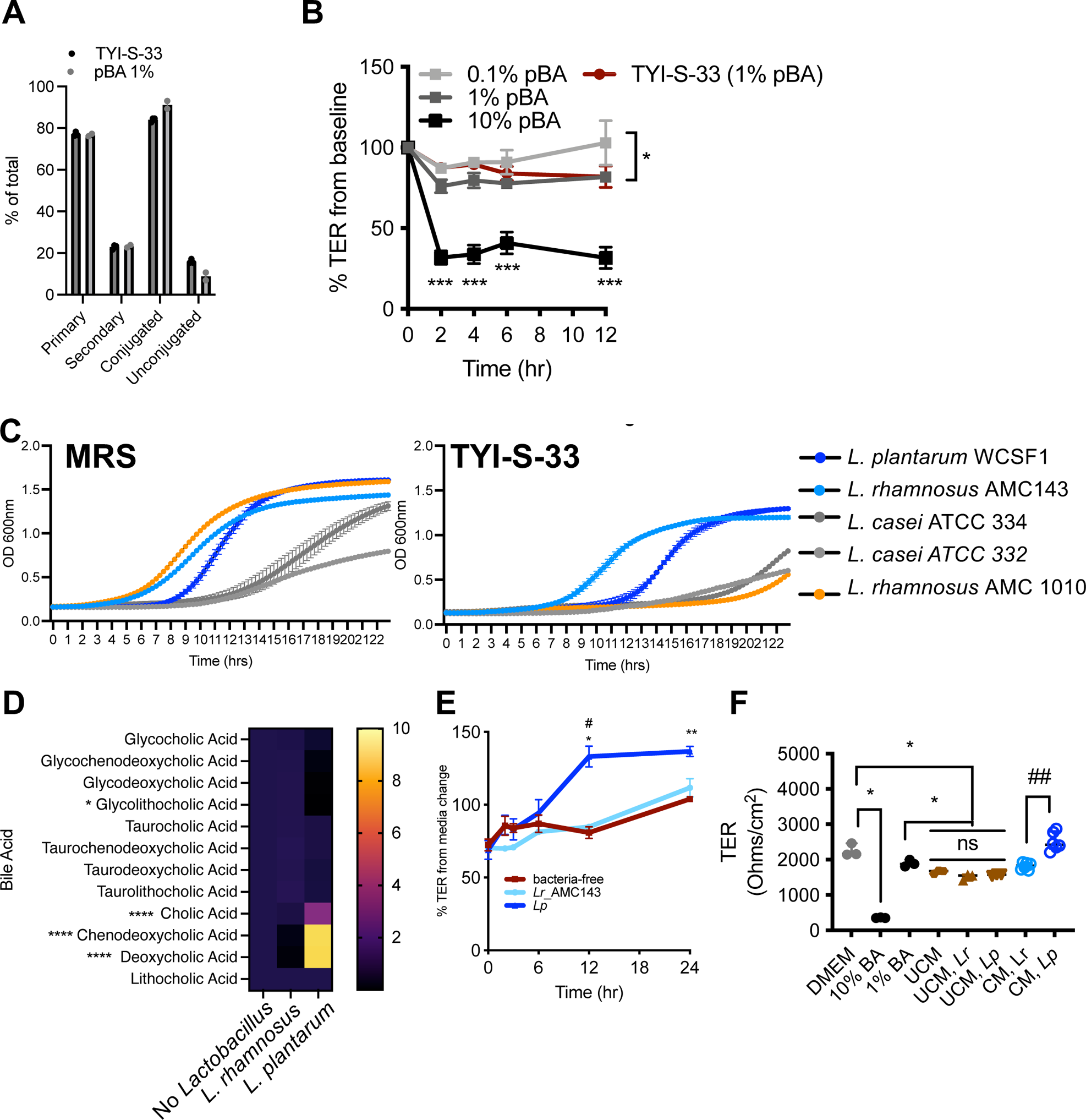
Physiological bile acids differentially support *Lactobacillus spp.* growth and protective functions of *Lactobacillus* on bile-acid induced intestinal epithelial cell barrier injury are strain specific. **(A)** Compositional analysis by percentage of total bile acids represented by primary, secondary, conjugated and unconjugated types present in physiological bile acids mixture when dissolved in DMEM (pBA 1%) or protozoan media (TYI-S-33). **(B)** Transepithelial cell electrical resistance (TER) as % change from baseline in T-84 monolayers exposed do different concentrations of pBA in DMEM (0.1, 1.0 and 10%) or TYI-S-33 media containing 0.1% pBA. *P<0.05, Two-Way ANOVA for 0.1% pBA vs either 1% pBA or TYI-S-33 media, ***P<0.001. Two-Way ANOVA with Tukey’s post-test analysis for multiple comparisons for 10% pBA vs 0.1% pBA (median ± IQR, N=3 wells group). **(C)** Growth curves of different *Lactobacillus* strains in MRS media (left) or TYI-S-33 (right). Shown are means of at least 6 technical replicates from each strain. For *L. plantarum* and *L.* rhamnosus_AMC143, curves are the mean of two separate biological replicates with at least technical six replicates each **(D)** Heat map of bile acid profiles in TYI-S-33 media after 24 hours of growth of *Lp* or *Lr*_AMC143 relative to baseline fresh media. **(E)** TER as % change from baseline in T-84 monolayers exposed to TYI-S-33 with either log-phase 10^6^ *Lp* or *Lr*_AMC143 or bacteria-free media. * P<0.05, ** P<0.01 for bacteria-free vs *Lp* and # P<0.05 for *Lp* vs *Lr*_AMC143, Two-Way ANOVA with Tukey’s post-test analysis for multiple comparisons (mean ± IQR, N=3 per group). **(F)** Baseline TER at 30 minutes after media change from DMEM to TYI-S-33 alone, or TYI-S-33 containing log-phase 10^6^ *Lp* or *Lr*_AMC143, or filter-sterilized conditioned TYI-S-33 wherein *Lp* or *Lr*_AMC143 were cultured for 24 hours, as indicated. *P<0.05 for DMEM vs 10% BA (Kruskal-Wallis with Dunn’s correction for multiple comparisons DMEM vs 10% BA or 1% BA), *P<0.05 for DMEM or 1% BA vs UCM and UCM*^Lr^* and UCM*^Lp^* (Kruskal-Wallis with Dunn’s test for multiple comparisons DMEM vs 1% BA, UCM, UCM*^Lr^*, and UCM*^Lp^*). ##P<0.01 for CM*^Lr^* vs CM*^Lp^* (Mann-Whitney U-test) (median ± IQR, N=3-6 per group).

We next examined whether the reported barrier-protective properties of *Lactobacillus* strains could protect from bile acid-induced disruptions in epithelial barrier integrity^13^. We first identified *Lactobacillus spp.* strains that were bile tolerant, as defined by achieving stationary phase growth in <24 hours in 1% BA-containing TYI-S-33 media (**Fig. 5C**). Two strains, *L. plantarum* WCSF-1 and *L. rhamnosus*^AMC143^ met these criteria. Both reach stationary phase at approximately similar times when grown in either MRS or TYI-S-33 media (**Supplemental Fig. 5**). Moreover, consistent with known differences in their functional *bsh* activity ^32,33^, both strains differed in their ability to deconjugate bile acids in vitro (**Fig. 5D**). Incubation with *bsh*-expressing *L. plantarum* WCSF-1 but not *L. rhamnosus*^AMC143^ protected T84 monolayers from pBA-induced barrier injury. The amelioration of bile acid-induced barrier defects occurred within 6 hours of incubation with viable *L. plantarum*WCSF-1 (**Fig. 5E**). Furthermore, this protective effect was also transferrable through application of filter-sterilized TYI-S-33 conditioned media (CM) within which *L. plantarum*^WCSF-1^ was grown overnight (**Fig. 5F**). Importantly, barrier protection was not observed with unconditioned media (UCM), or conditioned media from *L. rhamnosus*^AMC143^.

### *Giardia* directly antagonizes growth-promoting effects of commensal *Lactobacillus* strains in protein-deficient mice

Mono-colonization with select *L. plantarum* strains can maintain infant mouse growth despite undernutrition ^12^. To directly test the interactions between *Lactobacillus spp.* and *Giardia* during undernutrition, we used gnotobiotic techniques to selectively colonize germ-free (GF) mice fed a PD diet. Because 16S rRNA amplicon sequencing could not definitively confirm *Lactobacillus* species-level assignments, we designed a consortium of *Lactobacillus spp.* mixture (*L. spp. mix*) using published beneficial functions: *L. plantarum* known to improve intestinal barrier and promote growth; *L. casei* reported to diminish *Giardia* severity in a murine model ^26, 27^ and promoting expression of the intestinal transport protein PEPT1 ^21^*; L. johnsonii* a known antagonist of *Giardia* replication^17^; *L. rhamnosus* that regulates host responses to *Giardia* in a different murine model^16^. In these experiments, GF mice were weaned onto the PD diet in gnotobiotic isolators, prior to transfer into individually ventilated cages in the gnotobiotic barrier facility. First, compared with PBS-challenged GF controls, GF mice conventionalized with fecal intestinal microbiota (FMT) from PD-diet fed *Giardia*-free SPF mice demonstrated rapid weight loss and moribund condition requiring termination of the experiment by 1-week post-FMT (**Fig. 6A**). Whereas *Giardia* monoassociation had no direct effect on weight gain through up to 2 weeks post-challenge, selective colonization with *L. spp.* mix or *L. plantarum* alone promoted similar levels of weight gain with similar kinetics. The presence of *Giardia* antagonized the growth benefit from *L. spp. mix.* Fecal bile acid profiling confirmed the presence of functional bsh activity in *Giardia* + *L. spp. mix* co-colonized mice that was like *L. plantarum* alone, and not the result of *Giardia* activity, consistent with the current understanding that *Giardia* is incapable of bile salt deconjugation (**Supplemental Figure 6B**). Consequently, *Lactobacillus* significantly promoted weight gain during protein deprivation only when *Giardia* was absent (**Fig. 6B**). *Giardia* colonized the SI with similar efficiency in both mono-association and in combination with *L. spp. mix* (**Fig. 6C**). Although *L. johnsonii* has been shown to expedite *Giardia* clearance in nourished mice ^17^, this strain was not recovered from mice co-challenged with *L. johnsonii* in combination with *Giardia* during protein deficiency (**Supplemental Figure 6C**). *Lactobacillus spp.* duodenal and cecal colonization density as determined by colony counts on MRS agar were also similar regardless of *Giardia* exposure in *L. mix spp.* challenged mice (**Fig. 6D**).

**Figure 6.**
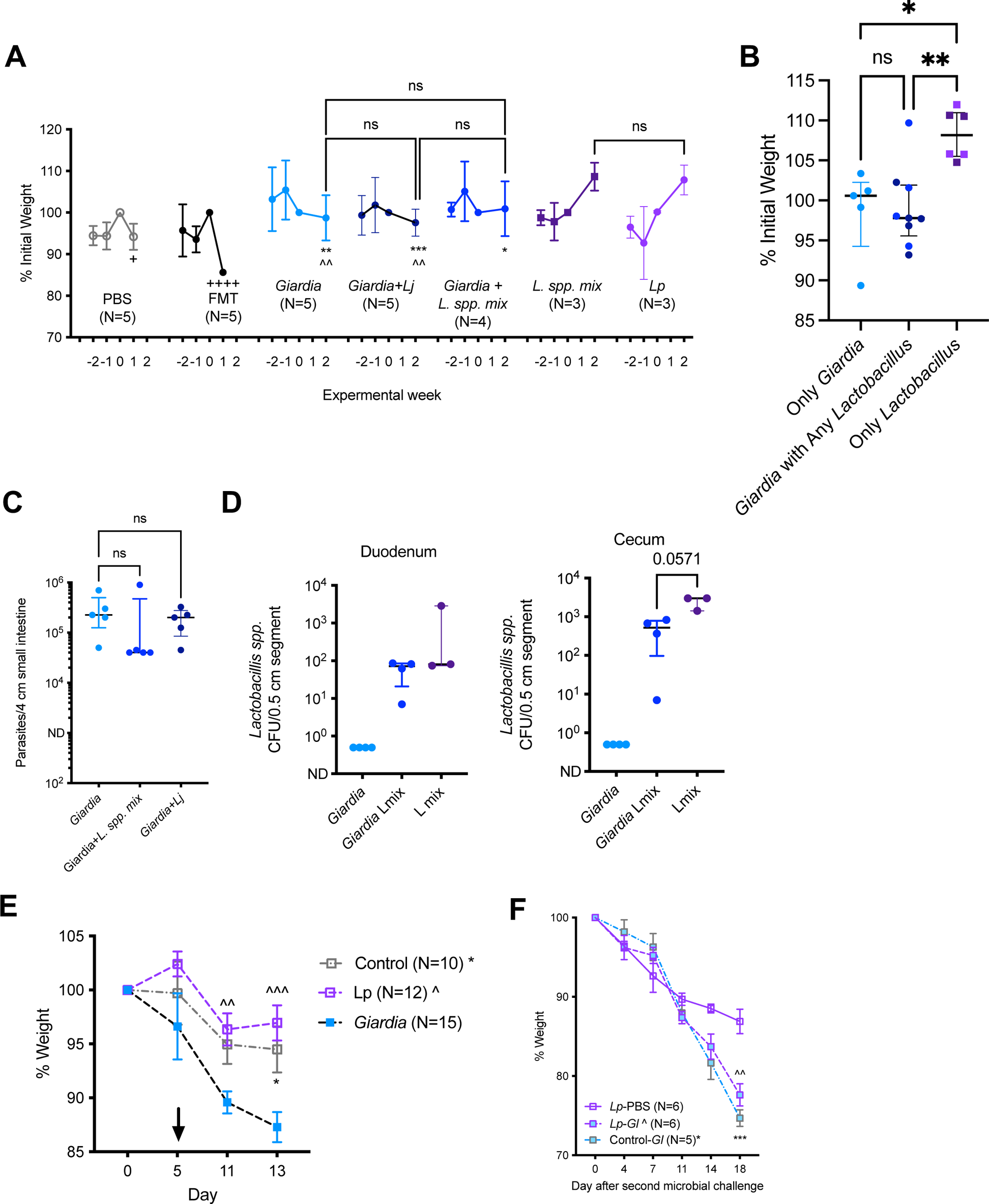
*Giardia* directly antagonizes growth-promoting *Lactobacillus spp.* in gnotobiotic mouse models of protein malnutrition. **(A**) Growth as % weight on the day of microbial challenge (0), beginning two weeks prior to and through 1 after challenge with PBS or fecal microbiota from SPF protein deficient mice (FMT) and two weeks after challenge with either 10^4^ axenic *G. lamblia* cysts or 10^6^ *Lactobacillus* strains alone or together as indicated. *L. spp. mix* = 10^6^ each of *Lj, Lp, Lr_*AMC010, *Lr_*AMC143, and *L. casei*. Three- to six-week-old germ-free (GF) mice were fed a protein deficient diet for the two weeks prior to transfer from isolators and oral gavage with indicated microbial challenge. Experiments were performed sequentially due to limited availability of age-matched GF mice. ^++++^P<0.0001 for PBS vs FMT (week 0-1); ^^^^P<0.01 *Lp* vs *Giardia* or *Giardia*+*Lj* (week 0-2); *P<0.05 for *L. spp. mix* vs *Giardia+ L. spp. mix,* **P<0.01 for *L. spp. mix vs Giardia,* ***P<0.001 for *L. spp. mix* vs *Giardia*+*Lj*, Two-Way ANOVA with Tukey’s post-test analysis for multiple comparisons (mean ± SEM, N=3-5 per group). **(B)** Growth as % initial weight through two weeks aggregated by presence or absence of *Giardia* challenge or any *Lactobacillus* strain. *P<0.05, **P<0.01 (Kruskal-Wallis with Dunn’s correction for multiple comparisons, median ± IQR, N=5-9 per group). **(C)** *Giardia* trophozoites in the small intestine in mono-associated mice and mice co-colonized with *Giardia* and *Lactobacillus* strains (Kruskal-Wallis with Dunn’s test for multiple comparisons, median ± IQR, N=5 per group). **(D)** *Lactobacillus* colony recovery on MRS plates from small intestine and cecal tissues in *Giardia* mono-associated mice, mice co-colonized with *Giardia*+*L. spp. mix,* and mice colonized with *L. spp. mix* without *Giardia* (Kruskal-Wallis with Dunn’s test for multiple comparisons, median ± IQR, N=3-5 per group). **(E)** Growth of 8-16 week old GF *Rag2^−/−^* mice as % initial weight following challenge with *Giardia* or *Lp* alone, or no microbial challenge (Control). Mice were transitioned from control to protein deficient diet on day 5 as indicated by the arrow. ^^P<0.01, ^^^P<0.001 *Lp* vs *Giardia* and *P<0.05 Control vs *Giardia*; Two-way ANOVA with Tukey’s post-test analysis for multiple comparisons (mean ± SEM, N=10-15 per group. **(F)** Growth of a subset of control (N=5) or *Lp*-mono-associated (N=12) *Rag2^−/−^* mice for 13 days as % change following secondary challenge with either *Giardia* or PBS as indicated. Mice remained on a protein deficient diet. ^^P<0.01 for *Lp-Gl* vs *Lp*-PBS and ***P<0.001 for Control-*Gl* vs *Lp*-PBS. Two-way ANOVA with Tukey’s post-test analysis for multiple comparisons (mean ± SEM, N=5-6 per group).

Finally, we examined the reciprocal interactions between *Giardia* and barrier-protective and growth promoting *L. plantarum* WCSF-1 (*Lp*) in our model of more severe protein-malnutrition in immunodeficient mice (*Rag2*^−/−^). In these studies, we selectively monoassociated GF immunodeficient mice for 5 days with either *Lp* or *Giardia* for 5 days; then, all mice were transitioned to the PD diet. Whereas *Giardia* mono-association led to expected weight loss through two weeks post-challenge ^9^, *Lp*-monoassociated mice maintained body weight, which was slightly higher than control, germ-free PD-diet fed *Rag2*^−/−^ mice (**Fig. 6E**). After 13 days of colonization, we then performed a heterologous challenge with the second microbe; i.e. *Lp*-monoassocoiated mice were challenged with *Giardia*, and *Giardia*-monoassociated mice were challenged with *Lp*. When *Lp* was introduced into *Giardia*-colonized mice, there was a slight rescue in body weight compared to mice colonized with *Giardia* alone; PD-fed immunodeficient mice colonized with *Lp* alone had the most stable body weight (**Fig. 6F**). Furthermore, progressive weight loss in PD-fed *Giardia*-monoassociated immunodeficient mice could not be rescued by introducing *Lp (***Supplemental Fig 6D**). Regardless of the order in which *Giardia* was introduced, it colonized the SI to similar levels; *Lp* had no effect on trophozoite density within the duodenum (**Supplemental Fig. 6E**). Similarly, Lp colony counts in both duodenum and colon were comparable regardless of the order in which Lp was introduced, i.e. prior to or after *Giardia* colonization (**Supplemental Fig. 6F**). This indicates that neither microbe exerts antimicrobial action per se against the primary colonizer. However, *Giardia* introduced in either order shifted the biogeographic distribution of *Lp* from the duodenum into the colon (**Supplemental Fig. 6G**). This displacement may have prevented *Lp* from exerting growth-promoting functions in protein-undernourished mice.

## DISCUSSION

In the small intestine, metabolic and nutrient homeostasis is coupled with functions of the local microbial community, whose disruption can result in a loss of mucosal immunoregulation, aberrant nutrient absorption, and perturbed metabolism ^18^, sometimes with exacerbated inflammation^5^. Emphasis on microbe-mediated contributions to these pathologies have largely focused on the presence of pathogenic invaders, or a detrimental shift in the balance of resident microbes that enhance expression of pathobiont properties. To date, it is unclear how the loss of commensal microbe functions that provide beneficial physiological and homeostatic support contributes to observed pathologies. Here, we report a new gnotobiotic approach to elucidate the potential consequences of displacement of beneficial small intestinal microbes by an environmental invader, *Giardia*. The presence of *Giardia* within the murine host diminished mucosal homeostasis, impaired host growth, and most importantly, counteracted the beneficial, growth-promoting effects exerted by *Lactobacillus spp.* in an undernourished, protein-deprived host.

We and others have previously reported using murine models that *Giardia* induces compositional or functional shifts in resident intestinal microbiota by altering microbe-host co-metabolic activity, increased virulence traits of pathobionts, and/or co-infecting bacteria^34,35,36,10^. Here, we report that the presence of *Giardia* may also diminish the abundance, intestinal distribution, and/or physiological functions of autochthonous lactic acid bacteria, even if other shifts in microbial community profiles were not observed by 16S rRNA amplicon sequencing. Importantly, our gnotobiotic experiments indicated that the loss of beneficial functions of *Lactobacillus spp.* during *Giardia* co-colonization were not simply due to parasite-induced competitive exclusion of all *Lactobacillus* strains—only modest reductions in colony counts were observed in the upper small intestine of co-colonized mice. It is therefore more likely that *Giardia* alters the microenvironment leading to changes in commensal bacterial adaptations. In this way *Giardia* may magnify the uncoupling of *Lactobacillus* from mucosal-associated micro-niches as has been reported in undernourished mice evidenced by diminished *Lactobacillus*-specific IgA^18^. *Giardia* has also been shown to alter expression and downstream functions of non-pathogenic *E. coli* strains^35^. Alternatively, as we have previously published, *Giardia*-mediated alterations in host responses to bacterial ligands^37^ may also dampen signaling pathways necessary for host responses to commensal bacteria.

Our work revealed significant alteration of *Lactobacillus spp*. in our *Giardia* model. *Lactobacillus spp*. are a closely studied genus of high interest in the field of probiotic interventions for malnutrition, and they have well-defined bile acid metabolizing properties^24, 25^. It is known that bile enriched for conjugated bile salts with higher hydrophobicity^38^ and phosphatidylcholine^28^ promotes *Giardia* replication. In contrast, *Lactobacillus spp.* that deconjugate bile acids inhibit *Giardia* growth *in vitro*^39^ and accelerate *Giardia* clearance in nourished hosts^17^. In light of these prior observations, we were surprised to find that *Lactobacillus spp.* inhibition of *Giardia* colonization was impeded in the context of a protein-deficient host. On the surface, this finding is counter to the improved anthropometric and histological responses reported in undernourished BALB/c *Giardia* models treated with *L. casei*^MTCC^ ^1423^ ^40^. However, closer examination of the growth trajectories reported therein (accounting for differences in baseline weights of mice) reveals only modest benefits of *L. casei* on growth during malnutrition, or as an adjuvant to renourishment ^27, 40^. The presence of *Giardia* further limits this benefit, and moreover, diminishes recovery of *L. casei* from fecal specimens by up to 4-logs, even if *Giardia* fecal cyst shedding was only modestly decreased. Despite the robust growth advantage conferred by the *Lactobacillus spp.* used in our selectively colonized, protein undernourished murine model, the additional presence of *Giardia* significantly antagonized this benefit. Further studies are required to determine whether *Giardia* similarly diminishes specific autochthonous *Lactobacillus spp.* strains in humans, especially in a diet-dependent manner. It will also be important to determine if *Giardia* similarly interferes with *Lactobacillus spp.* probiotics used for undernourished children. Indeed, current limitations in understanding of these principles may contribute to lackluster results from *Lactobacillus* probiotic interventions for undernutrition^41^. Based on the 16S rRNA amplicon sequencing data from our model, and a lack of robust colonization competition between *Giardia* and *Lactobacillus ssp*. throughout the gut, we predict that resolving these complex interactions in human studies will require high-resolution approaches for accurate taxonomic characterization of microbial communities. Moreover, metagenomic and transcriptomic characterization of diverse intestinal microenvironments will aid in uncovering the influence of *Giardia* on the functional and spatial localization of resident bacteria within micro-niches. These approaches may help resolve the mixed data generated using amplicon sequencing between children with or without *Giardia* infection ^42, 43^. Additionally, it is important to consider that the important interactions between *Giardia* and commensal bacteria may overlap in function but track to different taxonomic designations in humans compared with rodent models, further highlighting the importance of future functional studies.

*Giardia* associates with increased small intestinal permeability in human cohorts and in our murine model^9^. While *Giardia* may have direct effects on intestinal epithelial cell (IEC) barrier function through alterations in MLCK and ZO-1^44^, our data suggests that perturbations in the small intestinal microenvironment of protein undernutrition, such as overrepresentation of primary bile acids, may directly harm IECs when beneficial commensal functions are lost. Variable strain-level properties of *Lactobacillus* are important determinants in recovery from bile-acid inducible injury of intestinal epithelial cells. Physiological concentrations of primary bile acids caused *in vitro* barrier defects that were mitigated by *L. plantarum*^WCSF-1^ which encodes four functional bile salt hydrolase genes. In contrast, *L. rhamnosu*s^AMC143^, which encodes one non-expressed bile salt hydrolase gene was unable to ameliorate bile-induced barrier defects. Due to technical challenges in facile manipulation of their *bsh* genes, it remains unknown which one or more of the four active *bsh* enzymes of *L. plantarum*^WCSF-1^ impart barrier protective functions; it is also unclear if the presence of *bsh* activity was incidental to other potential IEC-supportive properties present in *L. plantarum*^WCSF-1^ that are absent in *L. rhamnosus*^AMC143^. We note that others have also found strain-selectivity in health-promoting *Lactobacillus* during undernutrition: for example, intraspecies differences were seen in the linear growth benefit resulting from *L. plantarum*^WJL^ but not *L. plantarum*^NIZ02877^ ^12^. Future studies to pinpoint the specific mechanisms through which *L. plantarum* and other *Lactobacillus spp.* promote IEC barrier integrity and bile acid regulation are relevant not only for *Giardia* enteropathy, but likely for additional pathologies of the upper small intestine, e.g. disrupted bile acid profiles in children with environmental enteric dysfunction (EED) ^30^. Our work highlights the utility of gnotobiotic techniques to determine which colonizing species of *Lactobacillus* and *Bifidobacterium* are most relevant for *Giardia* interactions.

In conclusion, we investigated how how small intestinal invaders interact with resident microbiota in the undernourished gut. We report that outcomes of mucosal and nutrient dysregulation and impaired host growth during *Giardia* infection may not only be due the presence of a pathogen but the absence of beneficial microbial functions that promote host health. While ongoing studies examining pathogen virulence factors and interventions to neutralize them remain important, our work underscores the importance of considering the entire scope of functional consequences resulting from disrupted small intestinal ecology in undernourished children. Further elucidating the conditions that facilitate colonization of beneficial microbiota in the small intestine and moreover sustain their functional resilience is likely a necessary consideration when devising optimized therapies and novel interventions that not only eliminate the harm, but also promote health.

## ACKNOWLEDGEMENTS

This work was supported by NIH/NIAID R01AI151214 (LAB) and Young Investigator Grants for Probiotics Research from the Global Probiotics Council (APB, LAB). The National Gnotobiotic Rodent Resource Center at UNC is supported by NIH P30DK034987.

## MATERIALS AND METHODS

### Mice

Mouse experiments were conducted in strict accordance with recommendations in the Guide for the Care and Use of Laboratory Animals of the National Institutes of Health. The mouse experimental protocol was approved by the Institutional Animal Care and Use Committee at the University of North Carolina at Chapel Hill (IACUC Protocol# 23-126). For specific pathogen free (SPF) mouse experiments, all experiments used male wild-type C57Bl/6 J mice obtained from Jackson Laboratories. These mice were age-matched at 21 days of life and at least 10 g prior to shipping. Mice were housed in light cycles of 12:12 (12 h light and 12 h dark), at the set temperature of 72 °F (± 3 °F) and a set humidity range of 30–70%. Mice were housed 1–2 mice/cage in individually ventilated cages (IVCs) in the Division of Comparative Medicine BSL2 isolation cubicle facility at UNC-CH. For experiments in germ-free (GF) mice, all experiments used both male and female C57Bl/6 J wild-type and *Rag2^−/−^* (C57Bl/6 J background) mice at 8–16 weeks-old obtained from the National Gnotobiotic Rodent Resource Center at UNC-CH. For each experiment, mice were sorted into age and sex-matched intervention and control groups and singly housed upon transfer from isolators to the SPF cubicle facility. Mouse weights were obtained using a battery-operated digital scale (Ohaus) with a precision of +/− 0.01 g. Fecal pellets were obtained as previously described ^9, 10, 19^.

### Mouse diets

Mice were fed either the Protein deficient (PD) diet (Envigo, TD.110200) or the isocaloric control (CD) diets (Envigo, TD.08678). For experiments in SPF mice, mice were acclimated immediately onto their respective diets upon arrival. For experiments in wild-type GF mice, mice were acclimated to the PD diet while in the isolator environment for 1-2 weeks prior to microbial challenges in the Division of Comparative Medicine BSL2 isolation cubicle facility at UNC-CH. For experiments in *Rag2^−/−^* GF mice, mice were transferred to the cubicle facility and given microbial challenge prior to transition to the PD diet five days later as indicated in **Figure 6**.

### *Giardia lamblia* cyst preparation and colonization

For experiments in SPF mice, ultra-purified *G. lamblia* (assemblage B, H3 cysts) were acquired from Waterborne, Inc and as previously described^9^. Cysts were used within 48 days of arrival and rinsed in PBS 3 times using centrifugation at 400-600 g x10 minutes. Cysts were counted by hemacytometer and diluted in PBS as necessary for an inoculum of 10^6^ cysts/mL. Mice were challenged with 100 μl by oral gavage for a challenge dose of 10^5^ cysts/mouse. For mono-association or direct co-colonization with *Lactobacillus* experiments, axenic *G. lamblia* (assemblage B, H3 cysts) were acquired from *Giardia* mono-associated *Rag2^−/−^* (C57Bl/6) propagators in the National Gnotobiotic Rodent Resource Center at UNC-CH. Cysts were prepared from fresh fecal homogenates using 400 μl PBS per fecal sample, followed by gravity sedimentation and dilution to 10^4^ cysts/100 μl. Mice were challenged with 100 μl each.

### *Lactobacillus* strains, *in vitro* growth curves, and *in vitro* permeability assays

The strains listed in Table 1 were acquired as indicated and stored in glycerol stocks according to manufacturer protocols: *L. plantarum* WCSF-1 (BAA-793), *L. johnsonii* (332), *L. casei* (334). *L. rhamnosus*_AMC143 and *L.rhamnosus*_AMC010 were obtained from glycerol-preserved archived strains in the UNC Microbiome Core after isolation as previously described^33^. De Man– Rogosa–Sharpe (MRS) media was obtained from Gibco and mixed as either broth or agar according to manufacturer instructions. TYI-S-33 media was made as previously described^19^, but modified to be antibiotic-free. Individual colonies isolated on MRS agar plates were used to inoculate liquid MRS or TYI-S-33 media. 200 ul of inoculated culture was placed into 96 well plates in triplicate. Automated optical density (OD) readings were measured every 15 minutes for 22.5 hours using a closed-system TECAN plate reader set at 37°C in ambient atmosphere with shaking at 225 RPM.

**Table 1:** List of strains and sources.

| Strain | Source |
| --- | --- |
| *L. plantarum* WCSF-1 (BAA-793)<sup>45</sup> | ATCC |
| *L. johnsonii* (332) | ATCC |
| *L. casei* (334) | ATCC |
| *L. rhamnosus*_AMC143 | UNC Microbiome Core<sup>33</sup> |
| *L.rhamnosus*_AMC010 | UNC Microbiome Core<sup>33</sup> |

### *Lactobacillus* growth for preparation for mouse colonization

Single colonies of each *Lactobacillus* strain were selected from MRS agar plates and grown overnight in MRS broth at 37°C under ambient atmosphere. Overnight cultures were backdiluted (1:100 dilution) in 10ml MRS broth and incubated to early log phase growth (OD 0.2-0.4) at 37°C under ambient atmosphere. Bacterial cells were harvested by centrifugation, washed with PBS, and resuspended in PBS at a concentration of 10^7^ cells/ml. Mice were then challenged with 100 μl each per oral gastric gavage for a challenge inoculum of 10^6^ bacterial cells per strain per mouse.

### Fecal microbiota transplant

To transfer fecal microbiota from SPF PD-fed uninfected mice, fresh fecal pellets were collected serially between 7 and 21 days on diet. The pellets were then snap-frozen and stored at −80 °C. On the day of challenge, frozen fecal pellets were pooled, thawed, diluted in PBS (10 mg/mL of PBS), and homogenized. Mice were inoculated with 100 *μ*l of the homogenate by orogastric gavage as previously described^9^.

### 16S bacterial profiling

DNA was extracted from stool as previously described^46^. Fecal 16S rRNA amplicon libraries were prepared using Illumina Nextera two-stage PCR library protocol. Briefly, the 515F-806R primer set containing Illumina adaptors were used to run a limited cycle PCR (25 cycles), after which the amplicons were cleaned up using Axygen Ampure PCR Cleanup Beads. Cleaned amplicons were then subject to indexing using Nextera barcodes in an 8-cycle PCR. Barcoded Libraries were cleaned up using Axygen Ampure PCR cleanup beads and quantified using PicoGreen DS DNA reagent. Libraries were pooled at equimolar concentration and subject to sequencing on Illumina MiSeq 2×250 platform. Sequencing reads were analyzed using DADA2 ^47^and QIIME2^48^. The forward reads were truncated to 220bp and denoised with DADA2. Chimera were removed using the pooled method. The amplicon sequence variants (ASV) were classified based on the SILVA database (release 138). The ASV abundance table were normalized as previously described ^49^ to correct for different sequencing depth across samples.

### Detection and quantification of intestinal microbes and host gene expression

#### Giardia

*Giardia* trophozoites were detected by light microscopy and enumerated with a hemocytometer as previously described ^9^. Briefly, a small intestinal fragment of 4 cm in length was taken 1 cm from the pyloric sphincter (duodenum) and then opened longitudinally, minced, and placed in 4 mL of ice-cold PBS for 30–45 min. Trophozoites were counted in 10 μl aliquots using a hemocytometer.

#### Lactobacillus

For culture detection and enumeration of *Lactobacillus* from gnotobiotic mice, the same segment that was used for *Giardia* detection was homogenized and then plated using serial dilutions on MRS agar. For colonic *Lactobacillus* culture detection and enumeration, a 4 cm in length segment of distal colon was similarly obtained from each mouse. The colon contents were rinsed with ice-cold PBS and the rinsed segment was minced and homogenized prior to plating on MRS agar using serial dilutions. Colony counts were obtained after 48 hour incubation in 37°C in ambient atmosphere.

#### DNA extraction from fecal and intestinal segments

DNA extractions for all fecal samples were performed using a modified version of Qiagen MagAttract Microbial DNA Kit on KingFisher Flex instrument as previously described. DNA was extracted from duodenum samples using a modified Qiagen QIAamp DNA Extraction kit including lysis performed in ATL Lysis Buffer with Proteinase K and AL buffer. Lysate was loaded onto silica column and washed with AW1 and AW2 buffers, prior to elution and quantification as previously described^33^.

#### Lactobacillus and Bifidobacterium qPCR

We performed targeted qPCR using previously reported primers ^50^ to quantify *Lactobacillus* and *Bifidobacterium* from fecal and duodenum DNA extracts. Equivalent amounts of genomic DNA was amplified in duplicate or triplicate in 10 µL reactions consisting of 5 µL PowerSYBR master mix, 1 µL template DNA, 1 µLof Primer mix at 1 µM concentration, and PCR-grade water for fecal samples and a 1:100 dilution of template DNA from duodenum samples to dilute interfering host DNA. Amplification was performed on QuantStudio Q6 qPCR instrument, and quality was confirmed by melt curve analysis. Abundance of *Lactobacillus* and *Bifidobacterium* was determined using the delta delta Ct method, with universal 16S V4 used for normalization.

#### Intestinal gene expression from mice

For host gene expression from mice, jejunal segments (2-4 cm in length) were obtained immediately distal to the duodenum segment used for trophozoite detection. RNA extraction and cDNA synthesis were performed using QIAgen kits per the manufacturer’s directions and performed qPCR on a Quantstudio RealTime PCR machine (ThermoFisher) using SYBR Green. qPCR was performed using SYBR Green Primers for *RegIII*γ**, *MMP7*, *IL-22, Pept1, 18S, FXR*^51^, and *TGR5*^52^ are listed in Table 2, and were purchased from Sigma. Gene expression was normalized to the housekeeping gene β-actin, except for FXR and TGR5 which were normalized to 18S gene. In both instances, subsequent quantification was completed by the delta delta method using PBS control mice as a reference.

**Table 2:**
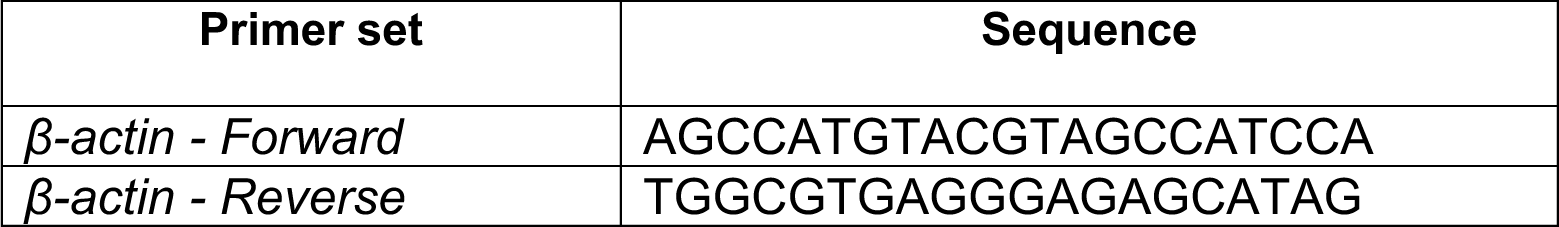
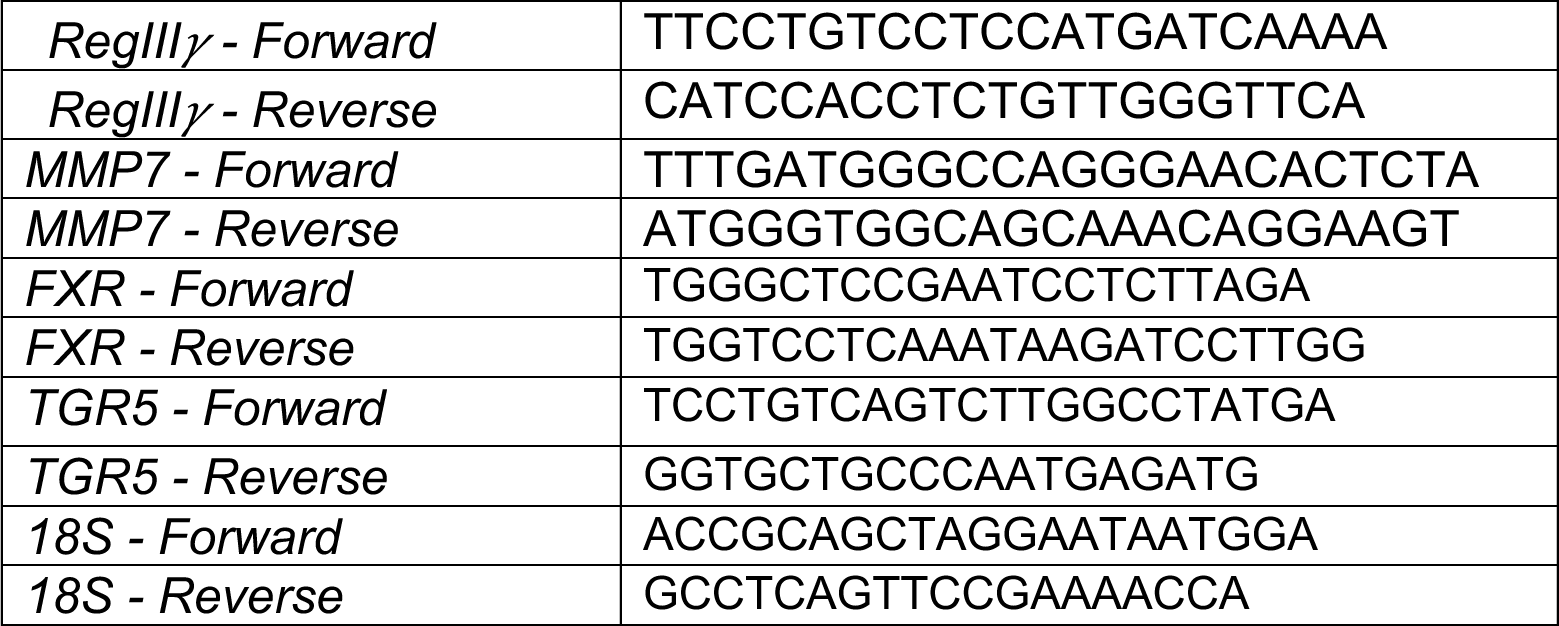
List of qPCR primers used to quantify host small intestinal gene expression.

### Crypt isolations and enumerations

Crypt isolations from mouse small intestine were performed as previously described^53^. Crypts were then separated from digested tissues using 100 μm cell strainers and counted using light microscopy in 10 μL droplets.

### Transepithelial cell resistance

**Mycoplasma-free T84** immortalized human carcinoma cells were obtained from ATCC (ATCC CCL-248). Confluent cells maintained in DMEM (Gibco) media supplemented with 10% heat inactivated fetal bovine serum (Sigma) and 1% penicillin-streptomycin mix; the latter was eliminated in experiments utilizing *Lactobacillus spp*. Cells were maintained in a water-jacketed incubator in a humidified environment maintained at 37°C with 5% CO_2_. To induce polarization, 2.5 E5 cells/ml were sown onto Transwell ® (3.0 uM, 12 mm polyester) inserts, and once cells adhered, transepithelial electrical resistance (TER) was serially measured using an EVOM2 epithelial voltohmmeter (World Precision Instruments) equipped with STX2 hand-held chopstick electrodes. TERs were measured on alternate days for 10-14 days, until their plateau at approximately 1200-2500 Ω/cm^2^; media was changed on alternate days by gently aspirating wells and replenishing with fresh pre-warmed media. Reagents for TEERS experiments consisted of TYI-S-33 media made as previously described^19^, DMEM with 0.1-10% physiologic bile acids (B8381, Sigma-Aldrich), or DMEM control media. For experiments described in Figure 5, serial TERs were measured for a total of 12 hours at 0.5-12 hour intervals. Data are represented as change from baseline and analyzed using 2-way ANOVA with post hoc Tukey test for multiple comparisons.

### Targeted amino acid and bile acid profiling

Blood was collected by cardiac puncture at the time of mouse necropsy, and serum was separated using BD (365967) serum separator additive microtainer tubes and centrifugation at 4000 x g rcf for 15 min. Fecal samples also collected at the time of sacrifice were snap-frozen in liquid nitrogen. Supernatants from TYI-S-33 media or DMEM enriched with physiologic bile acids were filter sterilized from bacteria-free or media containing *Lactobacillus* strains in 1 mL aliquots. All samples were stored at −80 °C and shipped on dry ice to the University of Pennsylvania Microbial Culture and Metabolomics Core. The Core performed targeted amino acids quantification using a Waters Acquity uPLC System with a Photodiode Array Detector. Amino acid concentrations are quantified via ultra-performance liquid chromatography (Waters Acquity UPLC system) with an AccQ-Tag Ultra C18 1.7 μm 2.1 x 100 mm column and a photodiode detector array. Analysis was performed using the UPLC AAA H-Class Application Kit (Waters Corporation, Milford, MA) according to manufacturer’s instructions. Standards are run at the beginning and end of each metabolomics run. Quality control checks (blanks and standards) are run every eight samples. Results are rejected if the standards deviate by greater than ± 5%. All chemicals and reagents are mass spectrometry grade. Samples without detectable analytes were assigned a value at half the limit of detection, which is 1 nmol/g in stool or 3 uM in plasma. For heatmaps, raw data was transformed to fold change relative to indicated reference groups. Where indicated, amino acids and bile acids were normalized to paired individual mouse intestinal permeability as determined by oral FITC-dextran (4.4 kDa gavage) as previously described ^9^. Simple linear regressions were used to correlate serum levels of bile acid and FITC-dextran.

### Statistics

For all murine experiments, animals were randomized into weight-matched groups at baseline. Due to the requirements to label infectious agents, the investigators were not blinded to allocation during experiments or to growth outcomes. Animal weight data were normalized to baseline absolute weights on day of challenge. For all growth assessments the absolute weights were transformed as percent of initial weight were analyzed using a two-way ANOVA test with repeated measures and Bonferroni post-test analysis for multiple comparisons. For comparisons between only two groups normalized data was compared using two-sided Unpaired t-tests and for non-normalized data two-sided Mann-Whitney U-test was used. The One-Way ANOVA with Holm-Sidak’s or Kruskal-Wallis with Dunn’s multiple comparisons tests were used for multi-group comparisons for normalized and non-normalized data, respectively. For 16S amplicon profiles, principal coordinates analysis (PCoA) was used to visualize the differences in microbial profiles at genus level between groups with the R package ‘vegan’. Ellipses indicate 95% confidence limits. PERMANOVA test (999 permutations) was used to analyze the differences in genus composition. Differences were considered significant at *P* < 0.05. Analyses for all data obtained from mouse models were analyzed with GraphPad Prism version 9.3. Data from all independent experiments are shown in the main figure or in the supplementary figures. Every datapoint indicates measured performed on independent biological replicates as either a single measurement or the mean of technical replicates. Simple linear regressions were used to perform correlation analyses in Figures 4D and Figure 6G. *Metabolomics methodology*: Serum and fecal amino acid and bile acid profiling data from CD-PBS and PD-PBS groups were filtered considering at least 50% of the valid values. Samples without detectable analyte values were replaced by the limit of detection (LoD) divided by the square root 2 (LoD of 3 uM for serum amino acids, 0.05 uM for serum bile acids, 0.5 nmol/g for faecal bile acids, and 1 nmol/g for faecal amino acids). Variables were selected by discrimination between the pair groups using Sparse Partial Least Squares Discriminant Analysis (sPLS-DA) and Principal Component Analysis (PCA) in MixOmics ^54^ on Rstudio. Statistical analyses were performed based on the normality test (Student’s t-test 10% FDR or Wilcoxon rank-sum Test 10% FDR) followed by a 1.5x fold change ratio for each comparison in both faecal and serum samples using “Volcano plot”, “ggplot2”, and “MixOmics” R packages. In the Volcano plot, red dots represent statistically significant overexpressed variables, while blue dots are the underrepresented variables in the first condition in the ratio of the paired comparison (e.g. PD-PBS/CD-PBS, PD-PBS is the first condition in the ratio).

---

**Supplemental Figure 1:**
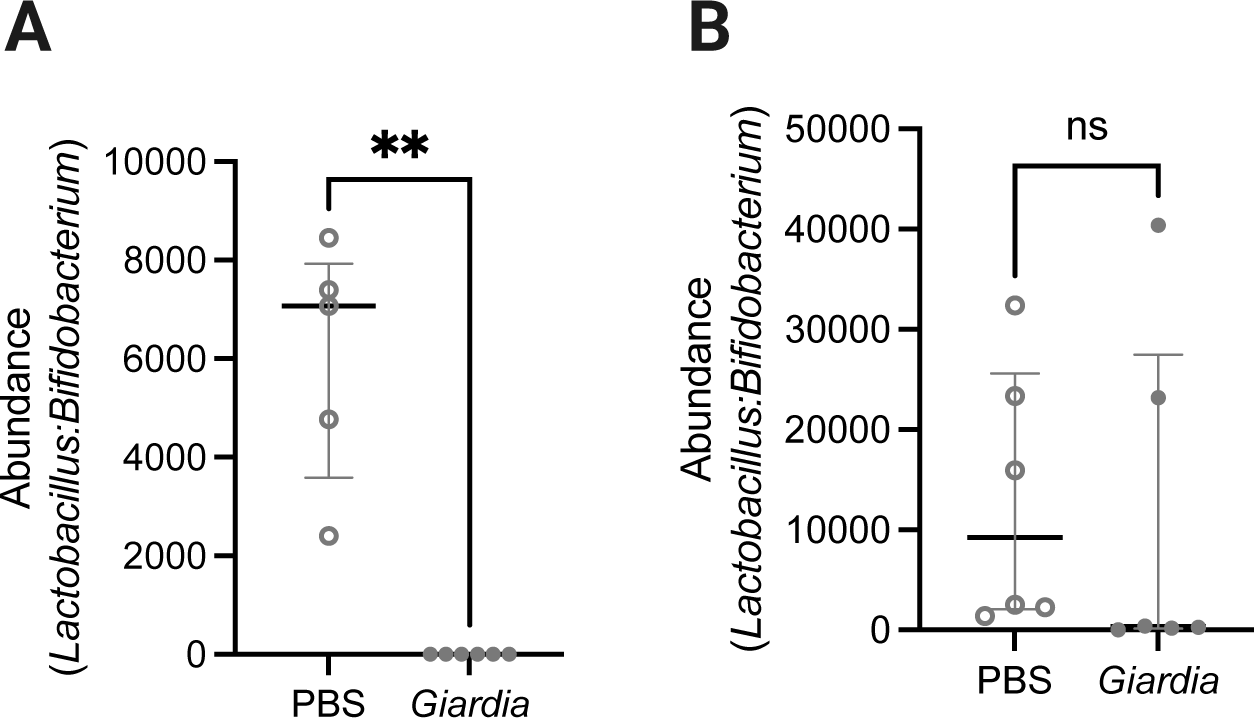
Ratios of abundances of *Lactobacillus* and *Bifidobacterium* in fecal samples from PD-fed, PBS- or *Giardia*-challenged mice from two separate experiments at Day 9 following *Giardia* challenge. (A) Significant difference in one experimental replicate: P<0.001 (Mann-Whitney U-test, median ± IQR, N=6 per group (B) Ratios tended to be lower with *Giardia,* but the difference was not statistically significant.

**Supplemental Figure 2:**
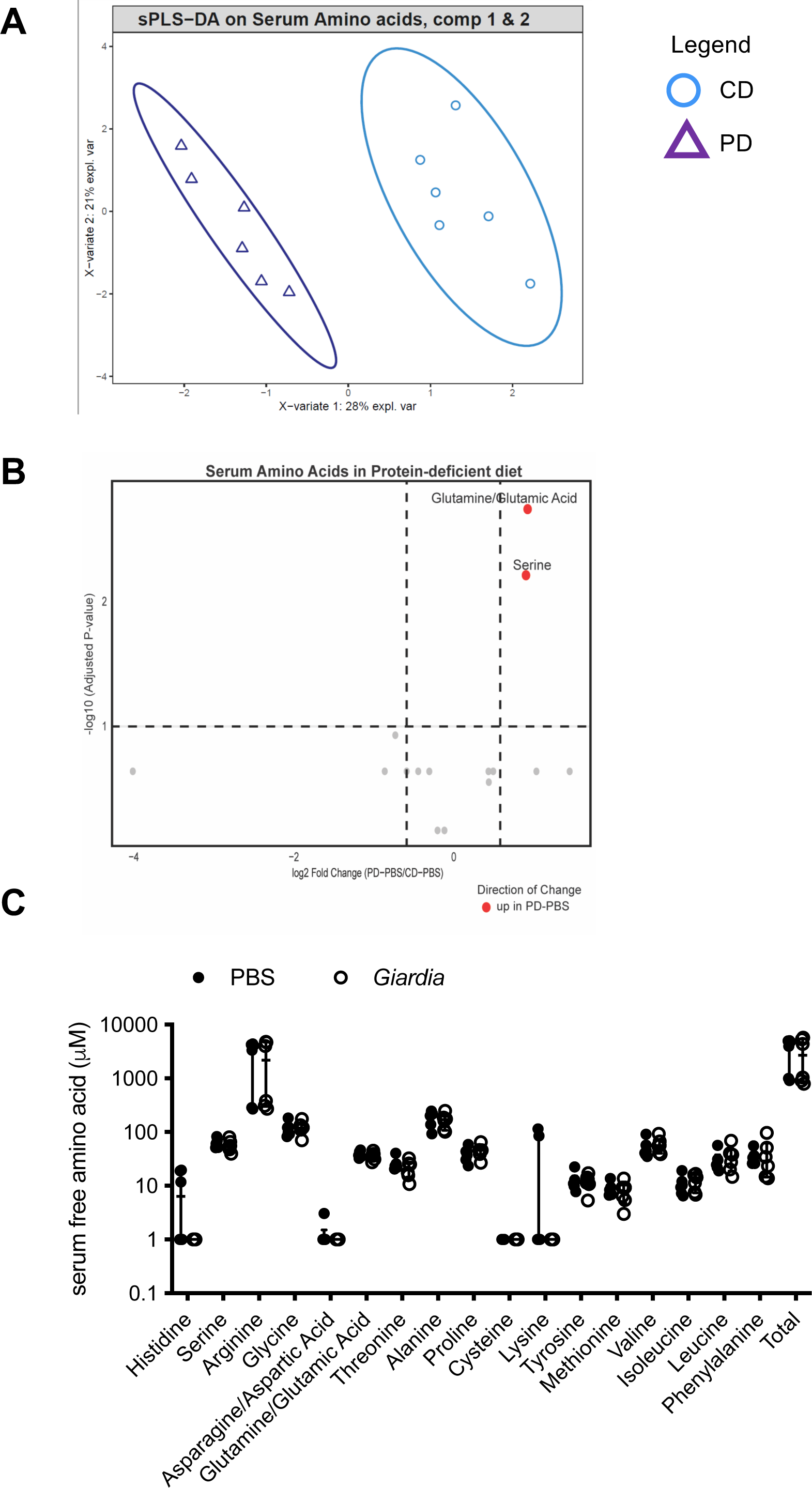
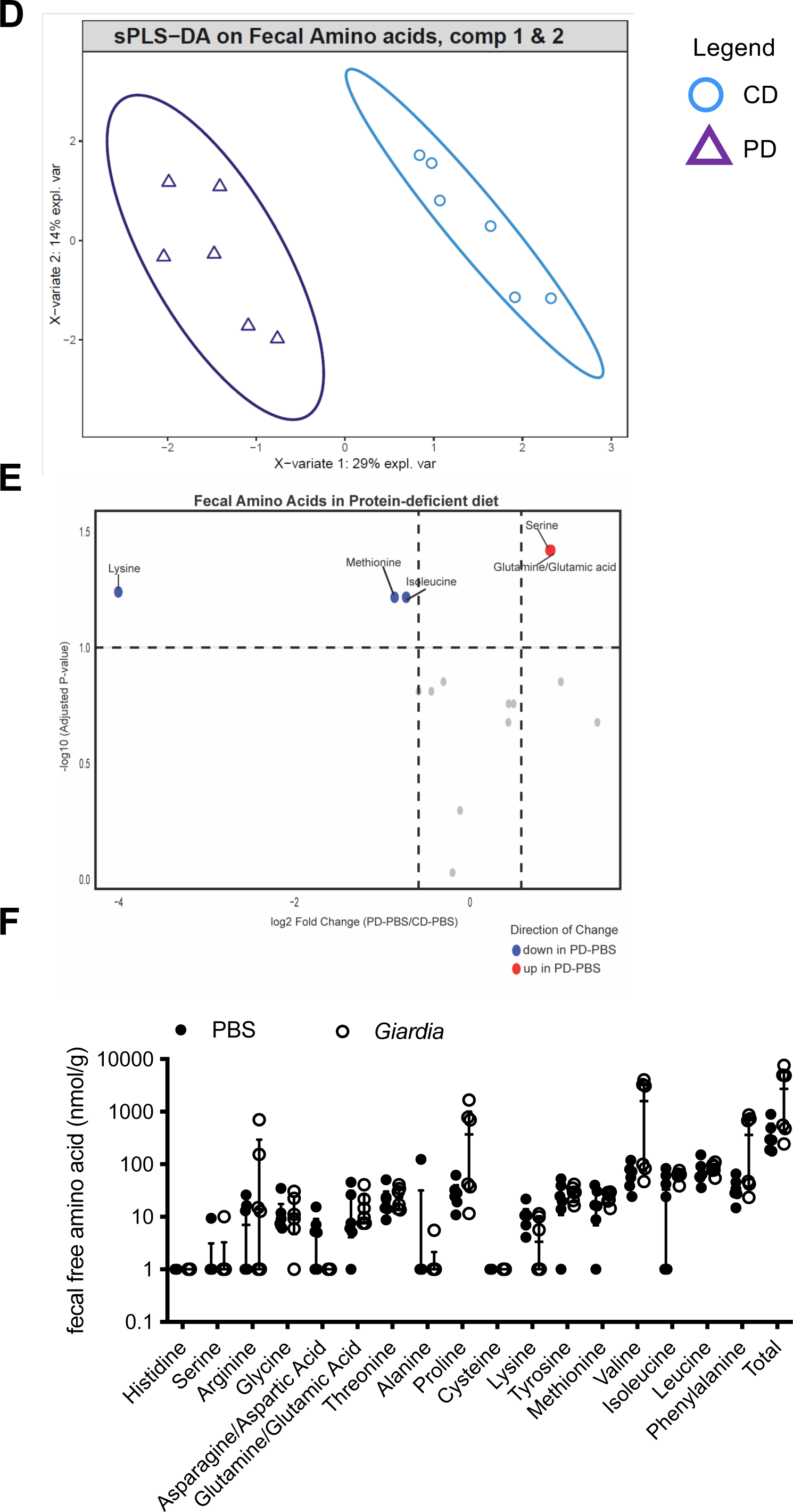
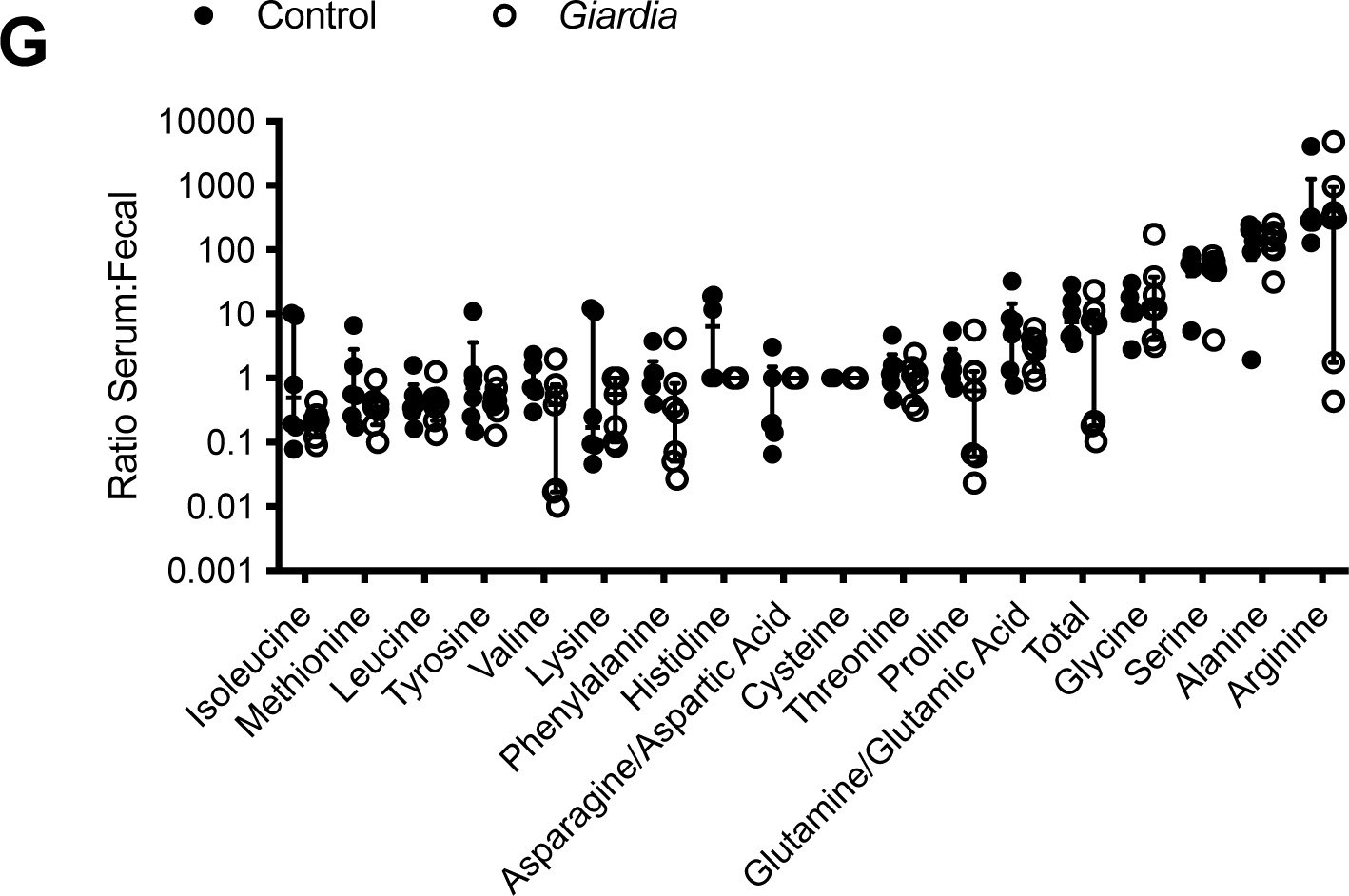
Diet is the primary driver of differences in serum and fecal amino acids. **(A):** Sample plots from sPLS-DA performed on the serum amino acids of CD-PBS and PD-PBS data showing a separation between them. **(B)** Serum amino acids showed significant differences during a protein-deficient diet (PD-PBS) compared to a normal diet (CD-PBS): high levels of serine and glutamine/glutamic acid (Wilcoxon rank-sum Test, 10% FDR). **(C)** concentrations of individual free amino acids in serum of PBS- or *Giardia*-challenged mice. **(D)** Fecal amino acids sample plots from CD-PBS and PD-PBS data (components 1 and 2) show the separation between the groups. **(E)** Volcano plot shows the modulation in the fecal amino acids in a protein-deficient diet: Lower amounts (blue dots) of Lysine, Methionine, and Isoleucine and higher amounts (red dots) of Serine and Glutamine/Glutamic acid (Wilcoxon rank-sum Test, 10% FDR). **(F)** concentrations of individual free amino acids in feces of PBS- or *Giardia*-challenged mice. **(G)** Ratios of serum:fecal amino acids in PBS- or *Giardia*-challenged mice.

**Supplemental Figure 3:**
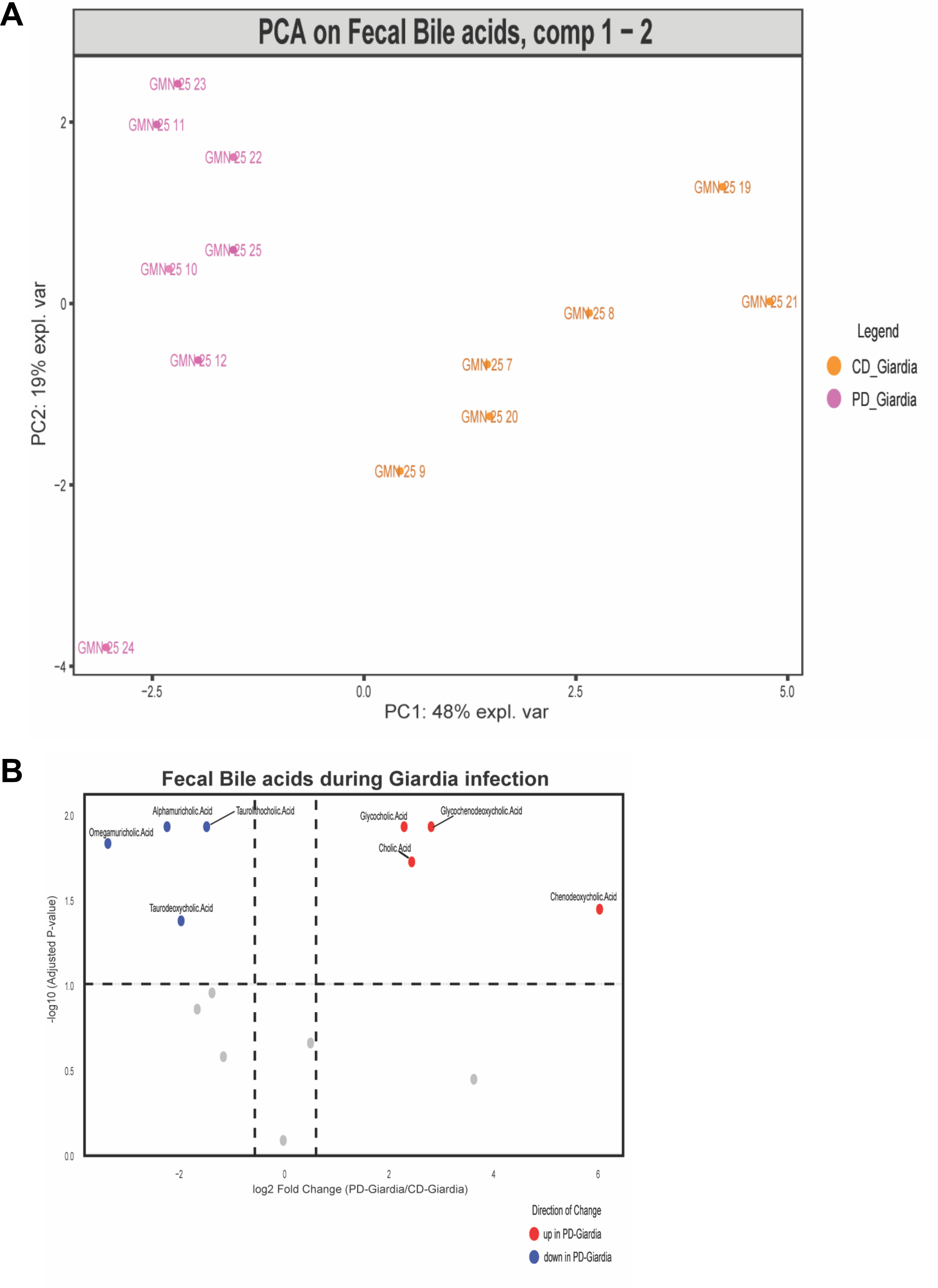
**(A)** PCA of fecal bile acids in PD-fed mice infected with *Giardia* (PD-Giardia) compared to mice infected during the control diet (CD-Giardia). **(B)** Volcano plot representing the modulation of the fecal bile acids in PD-Giardia: Lower amounts of Tauro-conjugated (Taurolithocholic acid, Taurodeoxycholic acid) and secondary bile acids (Omegamuricholic acid, Alphamuricholic acid, Taurolithocholic acid, Taurodeoxycholic acid) and higher amounts of primary (Cholic acid, Chenodeoxycholic acid) and glycol-conjguated (Glycocholic acid, Glycodenodeoxycholic acid) bile acids (Wilcoxon rank-sum Test, 10% FDR).

**Supplemental Figure 4:**
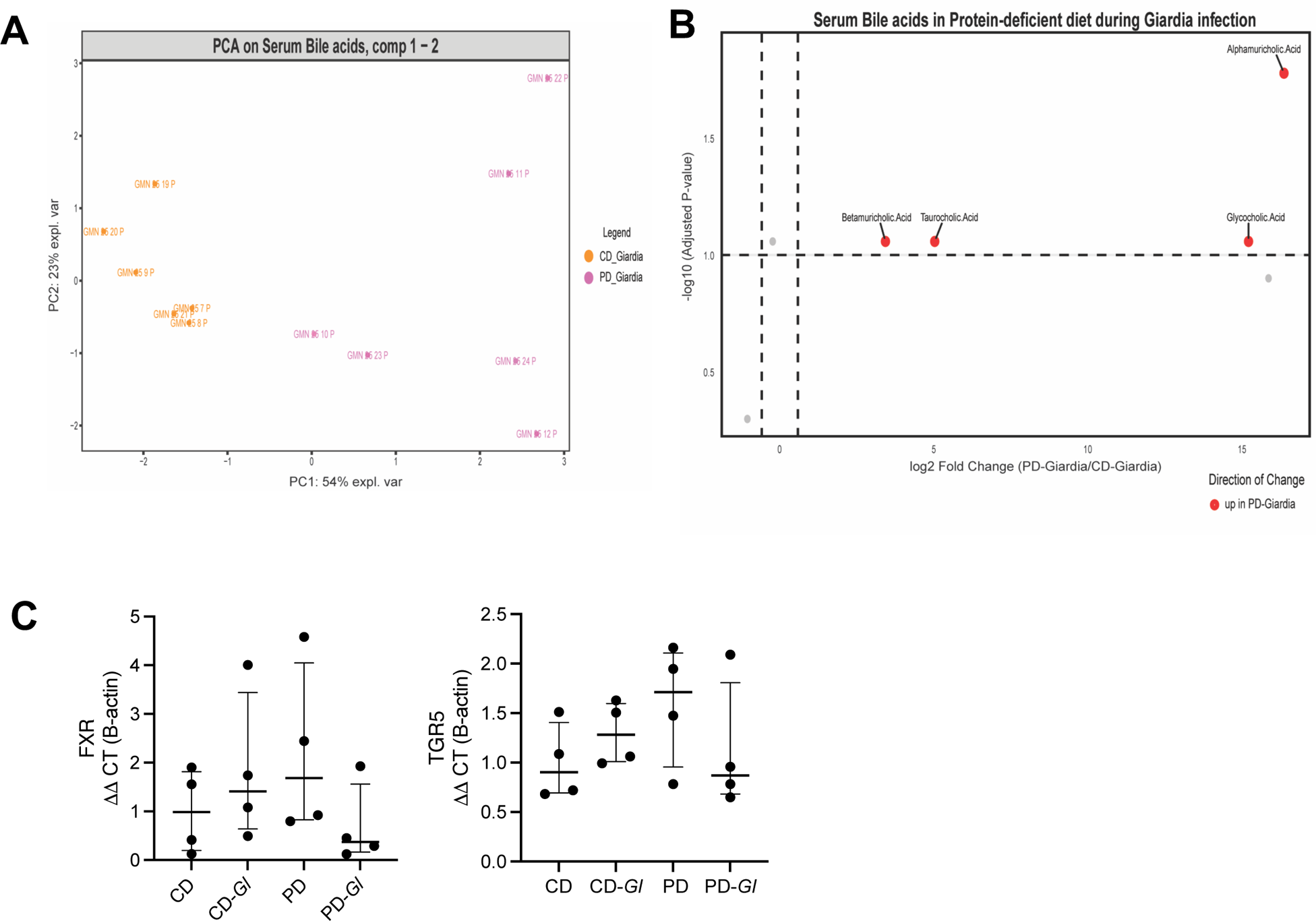
**(A)** PCA analysis of serum bile acids in PD-fed *Giardia*-infected mice compared to CD-fed *Giardia*-infected mice. **(B)** Volcano plot showing the significant abundance of serum bile acids in PD-fed and CD-fed *Giardia*-infected mice (Student’s t-test, 10% FDR). Red and blue dots are respectively the overrepresented and underrepresented bile acids in PD-*Giardia* compared to CD-*Giardia*. PD-Giardia showed higher levels of Tauro-conjugated (Taurocholic acid), glyco-conjugated (Glycocholic acid), and secondary (Betamuricholic acid) bile acids compared to CD-Giardia. **(C)** no differences observed between groups in bile acid homeostasis regulators *Fxr* or *Tgr5* in ileum tissue.

**Supplemental Figure 5:**
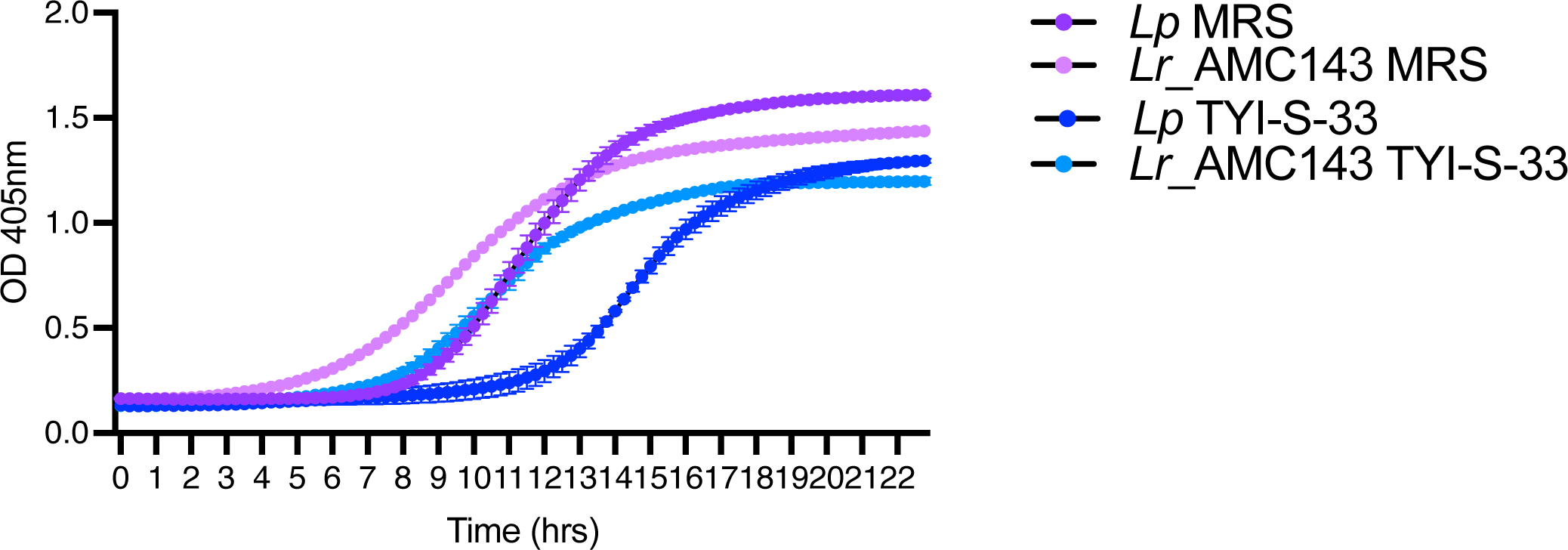
Comparison of growth rates of *L. plantarum* and *L. rhamonosus_*AMC143 in MRS and TYI-S-33 protozoa growth media.

**Supplemental Figure 6:**
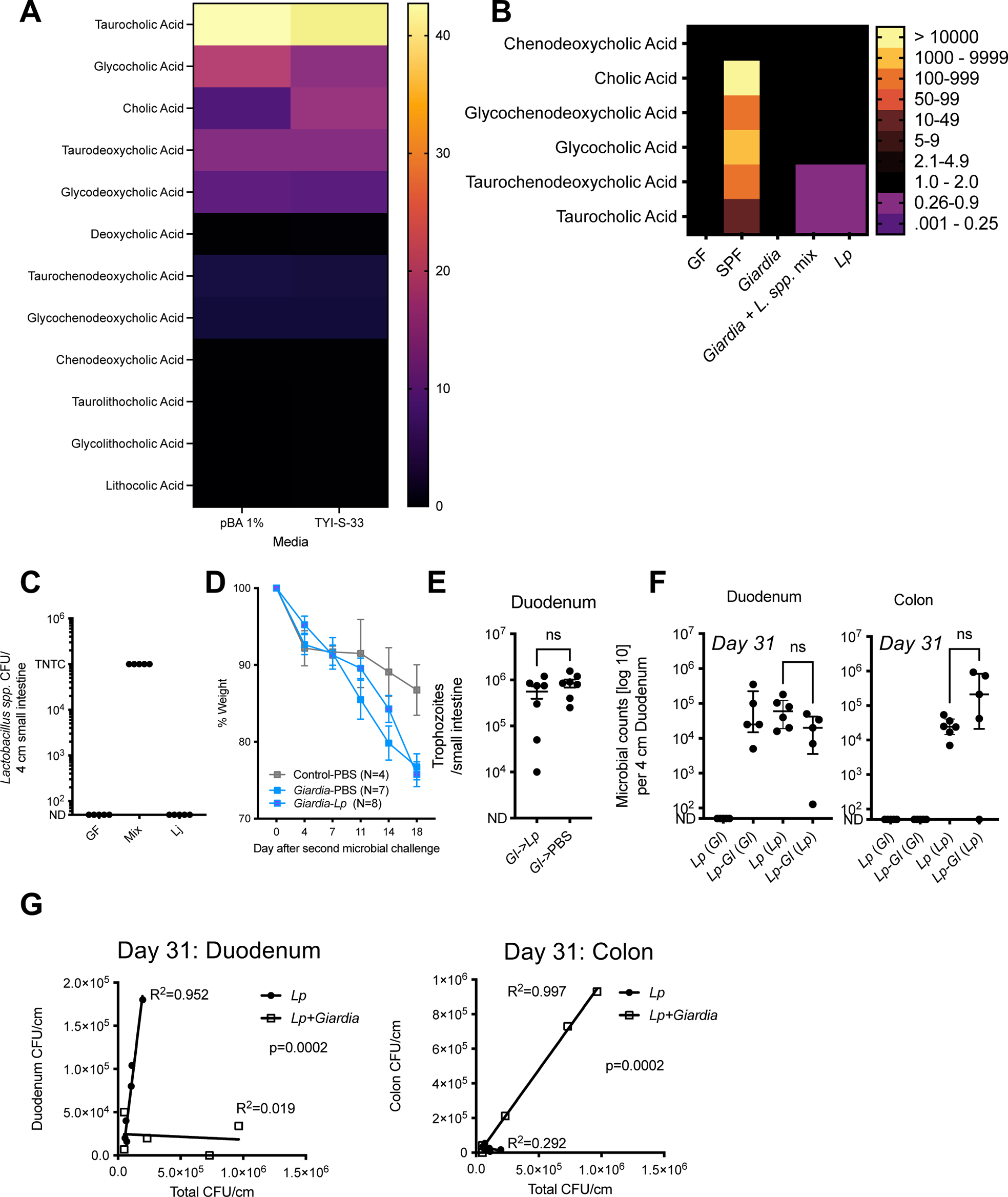
**(A)** quantification of indicated bile acids in 1% physiological bile acid mix (pBA 1%) or TYI-S-33 media. **(B)** quantification of indicated bile acids in stool of germ-free (GF) and SPF mice, and SPF mice colonized with either *Giardia* or *Lactobacillus spp.* mix alone, or with both microbes. **(C)** Quantification of *Lactobacillus spp.*, in small intestine segments of mice from Figure 5C. **(D)** change in body weight over time of immunodeficient mice following second microbe challenge. **(E)** Similar levels of *Giardia* trophozoites recovered from intestines of mice, regardless of order of microbe challenge. **(F)** No significant differences in the extent of *Lactobacillus* colonization regardless of order of microbe challenge. **(G)** *Giardia* challenge in *L. plantarum* colonized mice significantly correlates with displacement of *Lactobacillus* from the duodenum (left) to the colon (right), as revealed by simple linear regression. Duodenum: *Lp* R^2^=0.952, *Lp + Giardia* R^2^=0.019, p=0.0002; Colon: *Lp* R^2^=0.292, *Lp + Giardia* R^2^=0.997, p=0.0002.

## Notes

### Competing Interest Statement

The authors have declared no competing interest.

## REFERENCES

1. Roth DE, Krishna A, Leung M, Shi J, Bassani DG, Barros AJD. Early childhood linear growth faltering in low-income and middle-income countries as a whole-population condition: analysis of 179 Demographic and Health Surveys from 64 countries (1993-2015). Lancet Glob Health. 2017;5(12):e1249–e57. doi: 10.1016/S2214-109X(17)30418-7. PubMed PMID: 29132614; PMCID: PMC5695758.

2. Benjamin-Chung J, Mertens A, Colford JM, Jr., Hubbard AE, van der Laan MJ, Coyle J, Sofrygin O, Cai W, Nguyen A, Pokpongkiat NN, Djajadi S, Seth A, Jilek W, Jung E, Chung EO, Rosete S, Hejazi N, Malenica I, Li H, Hafen R, Subramoney V, Haggstrom J, Norman T, Brown KH, Christian P, Arnold BF, Ki Child Growth C. Early-childhood linear growth faltering in low- and middle-income countries. Nature. 2023;621(7979):550–7. Epub 20230913. doi: 10.1038/s41586-023-06418-5. PubMed PMID: 37704719; PMCID: PMC10511325.

3. Bartelt LA, Bolick DT, Guerrant RL. Disentangling Microbial Mediators of Malnutrition: Modeling Environmental Enteric Dysfunction. Cell Mol Gastroenterol Hepatol. 2019;7(3):692–707. Epub 20190107. doi: 10.1016/j.jcmgh.2018.12.006. PubMed PMID: 30630118; PMCID: PMC6477186.

4. Vonaesch P, Araujo JR, Gody JC, Mbecko JR, Sanke H, Andrianonimiadana L, Naharimanananirina T, Ningatoloum SN, Vondo SS, Gondje PB, Rodriguez-Pozo A, Rakotondrainipiana M, Kandou KJE, Nestoret A, Kapel N, Djorie SG, Finlay BB, Wegener Parfrey L, Collard JM, Randremanana RV, Sansonetti PJ, Afribiota I. Stunted children display ectopic small intestinal colonization by oral bacteria, which cause lipid malabsorption in experimental models. Proc Natl Acad Sci U S A. 2022;119(41):e2209589119. Epub 20221005. doi: 10.1073/pnas.2209589119. PubMed PMID: 36197997; PMCID: PMC9573096.

5. Chen RY, Kung VL, Das S, Hossain MS, Hibberd MC, Guruge J, Mahfuz M, Begum S, Rahman MM, Fahim SM, Gazi MA, Haque R, Sarker SA, Mazumder RN, Di Luccia B, Ahsan K, Kennedy E, Santiago-Borges J, Rodionov DA, Leyn SA, Osterman AL, Barratt MJ, Ahmed T, Gordon JI. Duodenal Microbiota in Stunted Undernourished Children with Enteropathy. N Engl J Med. 2020;383(4):321–33. doi: 10.1056/NEJMoa1916004. PubMed PMID: 32706533; PMCID: PMC7289524.

6. Vonaesch P, Morien E, Andrianonimiadana L, Sanke H, Mbecko JR, Huus KE, Naharimanananirina T, Gondje BP, Nigatoloum SN, Vondo SS, Kaleb Kandou JE, Randremanana R, Rakotondrainipiana M, Mazel F, Djorie SG, Gody JC, Finlay BB, Rubbo PA, Wegener Parfrey L, Collard JM, Sansonetti PJ, Afribiota I. Stunted childhood growth is associated with decompartmentalization of the gastrointestinal tract and overgrowth of oropharyngeal taxa. Proc Natl Acad Sci U S A. 2018;115(36):E8489-E98. Epub 20180820. doi: 10.1073/pnas.1806573115. PubMed PMID: 30126990; PMCID: PMC6130352.

7. Rogawski ET, Liu J, Platts-Mills JA, Kabir F, Lertsethtakarn P, Siguas M, Khan SS, Praharaj I, Murei A, Nshama R, Mujaga B, Havt A, Maciel IA, Operario DJ, Taniuchi M, Gratz J, Stroup SE, Roberts JH, Kalam A, Aziz F, Qureshi S, Islam MO, Sakpaisal P, Silapong S, Yori PP, Rajendiran R, Benny B, McGrath M, Seidman JC, Lang D, Gottlieb M, Guerrant RL, Lima AAM, Leite JP, Samie A, Bessong PO, Page N, Bodhidatta L, Mason C, Shrestha S, Kiwelu I, Mduma ER, Iqbal NT, Bhutta ZA, Ahmed T, Haque R, Kang G, Kosek MN, Houpt ER, Investigators M-EN. Use of quantitative molecular diagnostic methods to investigate the effect of enteropathogen infections on linear growth in children in low-resource settings: longitudinal analysis of results from the MAL-ED cohort study. Lancet Glob Health. 2018;6(12):e1319–e28. Epub 20181001. doi: 10.1016/S2214-109X(18)30351-6. PubMed PMID: 30287125; PMCID: PMC6227248.

8. Haberman Y, Iqbal NT, Ghandikota S, Mallawaarachchi I, Tzipi B, Dexheimer PJ, Rahman N, Hadar R, Sadiq K, Ahmad Z, Idress R, Iqbal J, Ahmed S, Hotwani A, Umrani F, Ehsan L, Medlock G, Syed S, Moskaluk C, Ma JZ, Jegga AG, Moore SR, Ali SA, Denson LA. Mucosal Genomics Implicate Lymphocyte Activation and Lipid Metabolism in Refractory Environmental Enteric Dysfunction. Gastroenterology. 2021;160(6):2055–71 e0. Epub 20210129. doi: 10.1053/j.gastro.2021.01.221. PubMed PMID: 33524399; PMCID: PMC8113748.

9. Giallourou N, Arnold J, McQuade ETR, Awoniyi M, Becket RVT, Walsh K, Herzog J, Gulati AS, Carroll IM, Montgomery S, Quintela PH, Faust AM, Singer SM, Fodor AA, Ahmad T, Mahfuz M, Mduma E, Walongo T, Guerrant RL, Balfour Sartor R, Swann JR, Kosek MN, Bartelt LA. Giardia hinders growth by disrupting nutrient metabolism independent of inflammatory enteropathy. Nat Commun. 2023;14(1):2840. Epub 20230518. doi: 10.1038/s41467-023-38363-2. PubMed PMID: 37202423; PMCID: PMC10195804.

10. Bartelt LA, Bolick DT, Mayneris-Perxachs J, Kolling GL, Medlock GL, Zaenker EI, Donowitz J, Thomas-Beckett RV, Rogala A, Carroll IM, Singer SM, Papin J, Swann JR, Guerrant RL. Cross-modulation of pathogen-specific pathways enhances malnutrition during enteric co-infection with Giardia lamblia and enteroaggregative Escherichia coli. PLoS Pathog. 2017;13(7):e1006471. Epub 20170727. doi: 10.1371/journal.ppat.1006471. PubMed PMID: 28750066; PMCID: PMC5549954.

11. Donowitz JR, Pu Z, Lin Y, Alam M, Ferdous T, Shama T, Taniuchi M, Islam MO, Kabir M, Nayak U, Faruque ASG, Haque R, Ma JZ, Petri WA, Jr. Small Intestine Bacterial Overgrowth in Bangladeshi Infants Is Associated With Growth Stunting in a Longitudinal Cohort. Am J Gastroenterol. 2022;117(1):167–75. doi: 10.14309/ajg.0000000000001535. PubMed PMID: 34693912; PMCID: PMC8715995.

12. Schwarzer M, Makki K, Storelli G, Machuca-Gayet I, Srutkova D, Hermanova P, Martino ME, Balmand S, Hudcovic T, Heddi A, Rieusset J, Kozakova H, Vidal H, Leulier F. Lactobacillus plantarum strain maintains growth of infant mice during chronic undernutrition. Science. 2016;351(6275):854–7. doi: 10.1126/science.aad8588. PubMed PMID: 26912894.

13. Yu Q, Yuan L, Deng J, Yang Q. Lactobacillus protects the integrity of intestinal epithelial barrier damaged by pathogenic bacteria. Front Cell Infect Microbiol. 2015;5:26. Epub 20150325. doi: 10.3389/fcimb.2015.00026. PubMed PMID: 25859435; PMCID: PMC4373387.

14. Wu H, Xie S, Miao J, Li Y, Wang Z, Wang M, Yu Q. Lactobacillus reuteri maintains intestinal epithelial regeneration and repairs damaged intestinal mucosa. Gut Microbes. 2020;11(4):997–1014. Epub 20200305. doi: 10.1080/19490976.2020.1734423. PubMed PMID: 32138622; PMCID: PMC7524370.

15. Goyal N, Rishi P, Shukla G. Lactobacillus rhamnosus GG antagonizes Giardia intestinalis induced oxidative stress and intestinal disaccharidases: an experimental study. World J Microbiol Biotechnol. 2013;29(6):1049–57. Epub 20130130. doi: 10.1007/s11274-013-1268-6. PubMed PMID: 23361971.

16. Goyal N, Shukla G. Probiotic Lactobacillus rhamnosus GG modulates the mucosal immune response in Giardia intestinalis-infected BALB/c mice. Dig Dis Sci. 2013;58(5):1218–25. Epub 20121221. doi: 10.1007/s10620-012-2503-y. PubMed PMID: 23263901.

17. Humen MA, De Antoni GL, Benyacoub J, Costas ME, Cardozo MI, Kozubsky L, Saudan KY, Boenzli-Bruand A, Blum S, Schiffrin EJ, Perez PF. Lactobacillus johnsonii La1 antagonizes Giardia intestinalis in vivo. Infect Immun. 2005;73(2):1265–9. doi: 10.1128/IAI.73.2.1265-1269.2005. PubMed PMID: 15664978; PMCID: PMC547090.

18. Huus KE, Bauer KC, Brown EM, Bozorgmehr T, Woodward SE, Serapio-Palacios A, Boutin RCT, Petersen C, Finlay BB. Commensal Bacteria Modulate Immunoglobulin A Binding in Response to Host Nutrition. Cell Host Microbe. 2020;27(6):909–21 e5. Epub 20200413. doi: 10.1016/j.chom.2020.03.012. PubMed PMID: 32289261.

19. Bartelt LA, Roche J, Kolling G, Bolick D, Noronha F, Naylor C, Hoffman P, Warren C, Singer S, Guerrant R. Persistent G. lamblia impairs growth in a murine malnutrition model. J Clin Invest. 2013;123(6):2672–84. doi: 10.1172/JCI67294. PubMed PMID: 23728173; PMCID: PMC3668820.

20. Qiu Y, Jiang Z, Hu S, Wang L, Ma X, Yang X. Lactobacillus plantarum Enhanced IL-22 Production in Natural Killer (NK) Cells That Protect the Integrity of Intestinal Epithelial Cell Barrier Damaged by Enterotoxigenic Escherichia coli. Int J Mol Sci. 2017;18(11). Epub 20171113. doi: 10.3390/ijms18112409. PubMed PMID: 29137183; PMCID: PMC5713377.

21. Neudeck BL, Loeb JM, Faith NG. Lactobacillus casei alters hPEPT1-mediated glycylsarcosine uptake in Caco-2 cells. J Nutr. 2004;134(5):1120–3. doi: 10.1093/jn/134.5.1120. PubMed PMID: 15113956.

22. Tako EA, Hassimi MF, Li E, Singer SM. Transcriptomic analysis of the host response to Giardia duodenalis infection reveals redundant mechanisms for parasite control. mBio. 2013;4(6):e00660–13. Epub 20131105. doi: 10.1128/mBio.00660-13. PubMed PMID: 24194537; PMCID: PMC3892777.

23. Ihara T, Tsujikawa T, Fujiyama Y, Bamba T. Regulation of PepT1 peptide transporter expression in the rat small intestine under malnourished conditions. Digestion. 2000;61(1):59–67. doi: 10.1159/000007736. PubMed PMID: 10671775.

24. O’Flaherty S, Briner Crawley A, Theriot CM, Barrangou R. The Lactobacillus Bile Salt Hydrolase Repertoire Reveals Niche-Specific Adaptation. mSphere. 2018;3(3). Epub 20180530. doi: 10.1128/mSphere.00140-18. PubMed PMID: 29848760; PMCID: PMC5976879.

25. Travers MA, Sow C, Zirah S, Deregnaucourt C, Chaouch S, Queiroz RM, Charneau S, Allain T, Florent I, Grellier P. Deconjugated Bile Salts Produced by Extracellular Bile-Salt Hydrolase-Like Activities from the Probiotic Lactobacillus johnsonii La1 Inhibit Giardia duodenalis In vitro Growth. Front Microbiol. 2016;7:1453. Epub 20160927. doi: 10.3389/fmicb.2016.01453. PubMed PMID: 27729900; PMCID: PMC5037171.

26. Shukla G, Singh S, Verma A. Oral Administration of the Probiotic Lactobacillus casei Ameliorates Gut Morphology and Physiology in Malnourished-Giardia intestinalis-Infected BALB/c Mice. ISRN Parasitol. 2013;2013:762638. Epub 20130921. doi: 10.5402/2013/762638. PubMed PMID: 27335861; PMCID: PMC4890930.

27. Shukla G, Sidhu RK. Lactobacillus casei as a probiotic in malnourished Giardia lamblia-infected mice: a biochemical and histopathological study. Can J Microbiol. 2011;57(2):127–35. doi: 10.1139/w10-110. PubMed PMID: 21326354.

28. Riba A, Hassani K, Walker A, van Best N, von Zezschwitz D, Anslinger T, Sillner N, Rosenhain S, Eibach D, Maiga-Ascofare O, Rolle-Kampczyk U, Basic M, Binz A, Mocek S, Sodeik B, Bauerfeind R, Mohs A, Trautwein C, Kiessling F, May J, Klingenspor M, Gremse F, Schmitt-Kopplin P, Bleich A, Torow N, von Bergen M, Hornef MW. Disturbed gut microbiota and bile homeostasis in Giardia-infected mice contributes to metabolic dysregulation and growth impairment. Sci Transl Med. 2020;12(565). doi: 10.1126/scitranslmed.aay7019. PubMed PMID: 33055245.

29. Raimondi F, Santoro P, Barone MV, Pappacoda S, Barretta ML, Nanayakkara M, Apicella C, Capasso L, Paludetto R. Bile acids modulate tight junction structure and barrier function of Caco-2 monolayers via EGFR activation. Am J Physiol Gastrointest Liver Physiol. 2008;294(4):G906–13. Epub 20080131. doi: 10.1152/ajpgi.00043.2007. PubMed PMID: 18239063.

30. Zhao X, Setchell KDR, Huang R, Mallawaarachchi I, Ehsan L, Dobrzykowski Iii E, Zhao J, Syed S, Ma JZ, Iqbal NT, Iqbal J, Sadiq K, Ahmed S, Haberman Y, Denson LA, Ali SA, Moore SR. Bile Acid Profiling Reveals Distinct Signatures in Undernourished Children with Environmental Enteric Dysfunction. J Nutr. 2021;151(12):3689–700. doi: 10.1093/jn/nxab321. PubMed PMID: 34718665; PMCID: PMC8643614.

31. Hegyi P, Maleth J, Walters JR, Hofmann AF, Keely SJ. Guts and Gall: Bile Acids in Regulation of Intestinal Epithelial Function in Health and Disease. Physiol Rev. 2018;98(4):1983–2023. doi: 10.1152/physrev.00054.2017. PubMed PMID: 30067158.

32. Lambert JM, Bongers RS, de Vos WM, Kleerebezem M. Functional analysis of four bile salt hydrolase and penicillin acylase family members in Lactobacillus plantarum WCFS1. Appl Environ Microbiol. 2008;74(15):4719–26. Epub 20080606. doi: 10.1128/AEM.00137-08. PubMed PMID: 18539794; PMCID: PMC2519332.

33. Arnold JW, Simpson JB, Roach J, Kwintkiewicz J, Azcarate-Peril MA. Intra-species Genomic and Physiological Variability Impact Stress Resistance in Strains of Probiotic Potential. Front Microbiol. 2018;9:242. Epub 20180220. doi: 10.3389/fmicb.2018.00242. PubMed PMID: 29515537; PMCID: PMC5826259.

34. Beatty JK, Akierman SV, Motta JP, Muise S, Workentine ML, Harrison JJ, Bhargava A, Beck PL, Rioux KP, McKnight GW, Wallace JL, Buret AG. Giardia duodenalis induces pathogenic dysbiosis of human intestinal microbiota biofilms. Int J Parasitol. 2017;47(6):311–26. Epub 20170222. doi: 10.1016/j.ijpara.2016.11.010. PubMed PMID: 28237889.

35. Gerbaba TK, Gupta P, Rioux K, Hansen D, Buret AG. Giardia duodenalis-induced alterations of commensal bacteria kill Caenorhabditis elegans: a new model to study microbial-microbial interactions in the gut. Am J Physiol Gastrointest Liver Physiol. 2015;308(6):G550–61. Epub 20150108. doi: 10.1152/ajpgi.00335.2014. PubMed PMID: 25573177; PMCID: PMC4360045.

36. Chen TL, Chen S, Wu HW, Lee TC, Lu YZ, Wu LL, Ni YH, Sun CH, Yu WH, Buret AG, Yu LC. Persistent gut barrier damage and commensal bacterial influx following eradication of Giardia infection in mice. Gut Pathog. 2013;5(1):26. Epub 20130830. doi: 10.1186/1757-4749-5-26. PubMed PMID: 23991642; PMCID: PMC3765889.

37. Burgess SL, Oka A, Liu B, Bolick DT, Oakland DN, Guerrant RL, Bartelt L. Intestinal parasitic infection alters bone marrow derived dendritic cell inflammatory cytokine production in response to bacterial endotoxin in a diet-dependent manner. PLoS Negl Trop Dis. 2019;13(7):e0007515. Epub 20190701. doi: 10.1371/journal.pntd.0007515. PubMed PMID: 31260452; PMCID: PMC6602177.

38. Farthing MJ, Keusch GT, Carey MC. Effects of bile and bile salts on growth and membrane lipid uptake by Giardia lamblia. Possible implications for pathogenesis of intestinal disease. J Clin Invest. 1985;76(5):1727–32. doi: 10.1172/JCI112162. PubMed PMID: 4056050; PMCID: PMC424194.

39. Perez PF, Minnaard J, Rouvet M, Knabenhans C, Brassart D, De Antoni GL, Schiffrin EJ. Inhibition of Giardia intestinalis by extracellular factors from Lactobacilli: an in vitro study. Appl Environ Microbiol. 2001;67(11):5037–42. doi: 10.1128/AEM.67.11.5037-5042.2001. PubMed PMID: 11679323; PMCID: PMC93268.

40. Shukla G, Sidhu RK, Verma A. Restoration of anthropometric, biochemical and histopathological alterations by Lactobacillus casei supplementation in Giardia intestinalis infected renourished BALB/c mice. Antonie Van Leeuwenhoek. 2012;102(1):61–72. Epub 20120302. doi: 10.1007/s10482-012-9713-3. PubMed PMID: 22382675.

41. Kamil RZ, Murdiati A, Juffrie M, Rahayu ES. Gut Microbiota Modulation of Moderate Undernutrition in Infants through Gummy Lactobacillus plantarum Dad-13 Consumption: A Randomized Double-Blind Controlled Trial. Nutrients. 2022;14(5). Epub 20220301. doi: 10.3390/nu14051049. PubMed PMID: 35268024; PMCID: PMC8912314.

42. Rouhani S, Griffin NW, Yori PP, Olortegui MP, Siguas Salas M, Rengifo Trigoso D, Moulton LH, Houpt ER, Barratt MJ, Kosek MN, Gordon JI. Gut Microbiota Features Associated With Campylobacter Burden and Postnatal Linear Growth Deficits in a Peruvian Birth Cohort. Clin Infect Dis. 2020;71(4):1000–7. doi: 10.1093/cid/ciz906. PubMed PMID: 31773126; PMCID: PMC7428392.

43. Mejia R, Damania A, Jeun R, Bryan PE, Vargas P, Juarez M, Cajal PS, Nasser J, Krolewiecki A, Lefoulon E, Long C, Drake E, Cimino RO, Slatko B. Impact of intestinal parasites on microbiota and cobalamin gene sequences: a pilot study. Parasit Vectors. 2020;13(1):200. Epub 20200419. doi: 10.1186/s13071-020-04073-7. PubMed PMID: 32306993; PMCID: PMC7168842.

44. Scott KG, Meddings JB, Kirk DR, Lees-Miller SP, Buret AG. Intestinal infection with Giardia spp. reduces epithelial barrier function in a myosin light chain kinase-dependent fashion. Gastroenterology. 2002;123(4):1179–90. doi: 10.1053/gast.2002.36002. PubMed PMID: 12360480.

45. van den Nieuwboer M, van Hemert S, Claassen E, de Vos WM. Lactobacillus plantarum WCFS1 and its host interaction: a dozen years after the genome. Microb Biotechnol. 2016;9(4):452–65. Epub 20160527. doi: 10.1111/1751-7915.12368. PubMed PMID: 27231133; PMCID: PMC4919987.

46. Arnold JW, Roach J, Fabela S, Moorfield E, Ding S, Blue E, Dagher S, Magness S, Tamayo R, Bruno-Barcena JM, Azcarate-Peril MA. The pleiotropic effects of prebiotic galacto-oligosaccharides on the aging gut. Microbiome. 2021;9(1):31. Epub 20210128. doi: 10.1186/s40168-020-00980-0. PubMed PMID: 33509277; PMCID: PMC7845053.

47. Callahan BJ, McMurdie PJ, Rosen MJ, Han AW, Johnson AJ, Holmes SP. DADA2: High-resolution sample inference from Illumina amplicon data. Nature methods. 2016;13(7):581–3. Epub 20160523. doi: 10.1038/nmeth.3869. PubMed PMID: 27214047; PMCID: PMC4927377.

48. Bolyen E, Rideout JR, Dillon MR, Bokulich NA, Abnet CC, Al-Ghalith GA, Alexander H, Alm EJ, Arumugam M, Asnicar F, Bai Y, Bisanz JE, Bittinger K, Brejnrod A, Brislawn CJ, Brown CT, Callahan BJ, Caraballo-Rodriguez AM, Chase J, Cope EK, Da Silva R, Diener C, Dorrestein PC, Douglas GM, Durall DM, Duvallet C, Edwardson CF, Ernst M, Estaki M, Fouquier J, Gauglitz JM, Gibbons SM, Gibson DL, Gonzalez A, Gorlick K, Guo J, Hillmann B, Holmes S, Holste H, Huttenhower C, Huttley GA, Janssen S, Jarmusch AK, Jiang L, Kaehler BD, Kang KB, Keefe CR, Keim P, Kelley ST, Knights D, Koester I, Kosciolek T, Kreps J, Langille MGI, Lee J, Ley R, Liu YX, Loftfield E, Lozupone C, Maher M, Marotz C, Martin BD, McDonald D, McIver LJ, Melnik AV, Metcalf JL, Morgan SC, Morton JT, Naimey AT, Navas-Molina JA, Nothias LF, Orchanian SB, Pearson T, Peoples SL, Petras D, Preuss ML, Pruesse E, Rasmussen LB, Rivers A, Robeson MS, 2nd, Rosenthal P, Segata N, Shaffer M, Shiffer A, Sinha R, Song SJ, Spear JR, Swafford AD, Thompson LR, Torres PJ, Trinh P, Tripathi A, Turnbaugh PJ, Ul-Hasan S, van der Hooft JJJ, Vargas F, Vazquez-Baeza Y, Vogtmann E, von Hippel M, Walters W, Wan Y, Wang M, Warren J, Weber KC, Williamson CHD, Willis AD, Xu ZZ, Zaneveld JR, Zhang Y, Zhu Q, Knight R, Caporaso JG. Reproducible, interactive, scalable and extensible microbiome data science using QIIME 2. Nat Biotechnol. 2019;37(8):852–7. doi: 10.1038/s41587-019-0209-9. PubMed PMID: 31341288; PMCID: PMC7015180.

49. Jones RB, Zhu X, Moan E, Murff HJ, Ness RM, Seidner DL, Sun S, Yu C, Dai Q, Fodor AA, Azcarate-Peril MA, Shrubsole MJ. Inter-niche and inter-individual variation in gut microbial community assessment using stool, rectal swab, and mucosal samples. Sci Rep. 2018;8(1):4139. Epub 20180307. doi: 10.1038/s41598-018-22408-4. PubMed PMID: 29515151; PMCID: PMC5841359.

50. Monteagudo-Mera A, Arthur JC, Jobin C, Keku T, Bruno-Barcena JM, Azcarate-Peril MA. High purity galacto-oligosaccharides enhance specific Bifidobacterium species and their metabolic activity in the mouse gut microbiome. Benef Microbes. 2016;7(2):247–64. Epub 20160203. doi: 10.3920/BM2015.0114. PubMed PMID: 26839072; PMCID: PMC4974821.

51. Kong B, Wang L, Chiang JY, Zhang Y, Klaassen CD, Guo GL. Mechanism of tissue-specific farnesoid X receptor in suppressing the expression of genes in bile-acid synthesis in mice. Hepatology. 2012;56(3):1034–43. Epub 20120712. doi: 10.1002/hep.25740. PubMed PMID: 22467244; PMCID: PMC3390456.

52. Velazquez-Villegas LA, Perino A, Lemos V, Zietak M, Nomura M, Pols TWH, Schoonjans K. TGR5 signalling promotes mitochondrial fission and beige remodelling of white adipose tissue. Nat Commun. 2018;9(1):245. Epub 20180116. doi: 10.1038/s41467-017-02068-0. PubMed PMID: 29339725; PMCID: PMC5770450.

53. Schoenborn AA, von Furstenberg RJ, Valsaraj S, Hussain FS, Stein M, Shanahan MT, Henning SJ, Gulati AS. The enteric microbiota regulates jejunal Paneth cell number and function without impacting intestinal stem cells. Gut Microbes. 2019;10(1):45–58. Epub 20180711. doi: 10.1080/19490976.2018.1474321. PubMed PMID: 29883265; PMCID: PMC6363071.

54. Le Cao KA, Boitard S, Besse P. Sparse PLS discriminant analysis: biologically relevant feature selection and graphical displays for multiclass problems. BMC Bioinformatics. 2011;12:253. Epub 20110622. doi: 10.1186/1471-2105-12-253. PubMed PMID: 21693065; PMCID: PMC3133555.

